# Antifungal therapeutic potential of *Candida albicans* Fun30: screening and validation of novel inhibitors against *Candida albicans*

**DOI:** 10.1101/2025.07.07.663444

**Authors:** Ketki Patne, Pragyan Parimita Rath, Faraz Mohd. Khan, R. Vijayan, Simran Sharma, Vijendra Arya, S. Gourinath, Rohini Muthuswami

## Abstract

WHO has listed *Candida albicans* as a fungal pathogen of priority. With the increasing incidence of *C. albicans* infection and limited treatment strategies, the identification of new drug targets and inhibitor molecules is of global concern. ATP-dependent chromatin remodelling proteins play a crucial role in regulating gene expression. Fun30 from *C. albicans* (*Ca*Fun30) mediates DNA end-resection during DNA double-strand break repair. It also transcriptionally co-regulates the expression of DNA damage response genes and genes involved in white-opaque switching. The gene is not essential for viability; however, the null mutant shows increased sensitivity to genotoxic agents. We hypothesised that the protein could be a potential therapeutic target against *C. albicans*. The structural model of *Ca*Fun30 was generated using homology modelling. Screening of a small molecule library identified 10 potential molecules, of which F1853-0039 was found as the lead inhibitor candidate. *In vitro* studies showed that the molecule had an MIC in the micromolar range against *C. albicans* with minimal effect on human THP-1 cells. Finally, coculture experiments showed that the inhibitor molecule prevented the formation of hyphae of *C. albicans* upon infecting THP-1 cells leading us to conclude that *Ca*Fun30 could potentially be developed as a therapeutic target against *C. albicans*.

## INTRODUCTION

Candidiasis is one of the most common fungal infections caused by the yeast *Candida*. More than 15 distinct *Candida* species, including *Candida albicans,* can cause disease. The organism grows as normal flora in the gastrointestinal tract and genitourinary system in healthy humans. However, overgrowth may occur due to an impaired immune system that leads to Candidiasis, which can also lead to fatal conditions such as candidemia. Candidemia is the most common fungal bloodstream infection in hospitalised patients.

The choice of drugs is limited to four classes of compounds: azoles, allylamines (terbinafine), polyenes (amphotericin B), pyrimidines (flucytosine), and echinocandins, of which azoles are the most widely used [1]. With an increased incidence of resistance, it is essential to identify newer drug targets.

Epigenetic modulators have emerged as promising drug targets [2]. In the case of *Candida* spp., histone deacetylase (HDAC) inhibitors have shown the potential to enhance the effect of azoles and to limit the emergence of antifungal resistance [3]. Therefore, we decided to investigate whether ATP-dependent chromatin remodelling proteins can also be potential drug targets.

ATP-dependent chromatin remodelling proteins regulate gene expression, replication, and DNA repair in eukaryotic organisms [4]. These proteins harness the energy released from ATP hydrolysis to reposition and/or evict nucleosomes, as well as mediate the exchange of histone variants [5]. The ATP-dependent chromatin remodellers are further divided into subfamilies, including Snf2 and Etl1 subfamilies. Fun30 is an ATP-dependent chromatin remodelling protein that belongs to the Etl1 subfamily. The protein was originally identified by a genetic screening in *Saccharomyces cerevisiae* [6], with homologs in almost all organisms. Fft1, 2, and 3 in *Schizosaccharomyces pombe*, Etl1 in mice, and SMARCAD1 in humans are the identified homologs [7,8]. All these proteins contain a conserved ATPase domain and a CUE domain. Fun30 from *S. cerevisiae* is a homodimer possessing DNA-stimulated ATPase activity [9]. The primary function of this protein is in the end resection step of the DNA during DNA double-strand break repair [8,10]. In addition, the protein also co-regulates gene expression and splicing [7,11–14].

The role of Fun30 in *C. albicans* (*Ca*Fun30) has recently been characterised. The protein is not essential for viability but mediates the response of the organism to genotoxic stressors and white-to-opaque switching [15,16]. Importantly, the protein regulates the expression of *RTT109*, which encodes for the fungal-specific histone acetyltransferase Rtt109 that mediates H3K56 acetylation [15]. Rtt109-mediated H3K56 acetylation is required for DNA damage response and, thus, pathogenesis in *C. albicans* [17]. Further, studies have shown that Rtt109-mediated H3K56ac modulates the transcription of *GPI15* and *GPI19* in *C. albicans* [18]. These two genes encode for proteins involved in the first step of GPI anchor biosynthesis. Thus, null mutants of *RTT109* have reduced GPI-APs at the cell surface, attenuated virulence, and increased susceptibility to azoles [18]. Inhibitors for Rtt109 are being developed [19]; however, they do not affect *C. albicans*’ growth.

As Fun30 regulates critical pathways, including white-to-opaque switching and hyphal morphogenesis, in *C. albicans*, we hypothesised that the protein could be a potential target to develop inhibitors against the pathogen.

Herein, we report the identification of an inhibitor of *Ca*Fun30. Ten potential ligands were identified by screening compounds from the database of Life Chemicals [20] using molecular docking tools like GOLD [21] and AutoDock 4.2 [22]. *In silico* and *in vitro* binding studies revealed three top compounds as good binding partners of the target protein *Ca*Fun30. Of these, F1853-0039 showed significant antifungal activity *in vitro,* suggesting that the molecule can be a promising candidate for developing a therapeutic regime.

## MATERIALS AND METHODS

### Chemicals

All the chemicals used in this study were of analytical grade and purchased from Sigma Aldrich (USA), Merck (India), SRL (India), and Fisher Scientific (USA). Restriction endonuclease enzymes were purchased from New England Biolabs (USA). Plasmid extraction kit and Gel Extraction kit were purchased from Qiagen (USA) and Thermo Scientific (USA). DNA and protein molecular weight markers were purchased from GeneDireX, Inc. and Thermofisher Scientific (USA). Bradford reagent was purchased from Sigma-Aldrich (USA). Ni^2+^-NTA sepharose resins were purchased from GE Healthcare (USA). Chemical compounds for inhibitor screening were purchased from Life Chemicals, Canada.

### Vectors and strains

The TA cloning vector (pTZ57R/T) was purchased from MBI Fermentas, USA and the expression vector pCold I was purchased from Addgene, USA. The bacterial strain DH5α (Bangalore Genei) was used for cloning, and BL21 (DE3) Rosetta cells (Merck) were used for expression studies. The SN152 strain was a kind gift from Prof. K. Natarajan, JNU, and the BWP17 strain was a kind gift from Prof. S.S. Komath, JNU.

### Cell culture

The bacterial strains were cultured in Luria-Bertani medium. *C. albicans* strains BWP17 and SN152 were cultured in Yeast Extract-Peptone-Dextrose (YEPD) media. THP-1 cells were cultured in Roswell Park Memorial Institute (RPMI 1640) medium. The three matched pair isolates of *C. albicans* and the drug-resistant strains of *C. auris* were a kind gift from Prof. Rajendra Prasad, Amity University.

### Structure prediction of *Ca*Fun30-c334

As the X-ray crystal structure of *Ca*Fun30 is unavailable, model building was done using homology modelling. The template sequence for homology modelling was searched by using the *Ca*Fun30 protein sequence as the query in the NCBI protein BLAST server (https://blast.ncbi.nlm.nih.gov/Blast.cgi) against the Protein Data Bank (<u>http://</u>www.rcsb.org/pdb). Multiple sequence alignment was performed by MUSCLE [23] and visualised using ESPript 3.0 [24] (https://espript.ibcp.fr/ESPript/ESPript/). Based on higher percentage identity and lower e-value in PDB blast results, the crystal structure of C-terminal region of Fun30 ATPase domain (PDB: 5GN1) of *S. cerevisiae* [25] was selected as template to predict the three-dimensional structure of the homologous region of *Ca*Fun30 which was named as *Ca*Fun30- c334 (Fig. 1A) using the Modeller stand-alone software [26] (https://salilab.org/modeller/). The structure was prepared by the Protein Preparation Wizard of Schrödinger [27,28]. The 3D structure of the modelled protein was visualised using PyMOL [29] and Discovery Studio [30]. PROCHECK [31] was used to validate the stereochemical structural quality of the predicted model.

**Figure 1.**
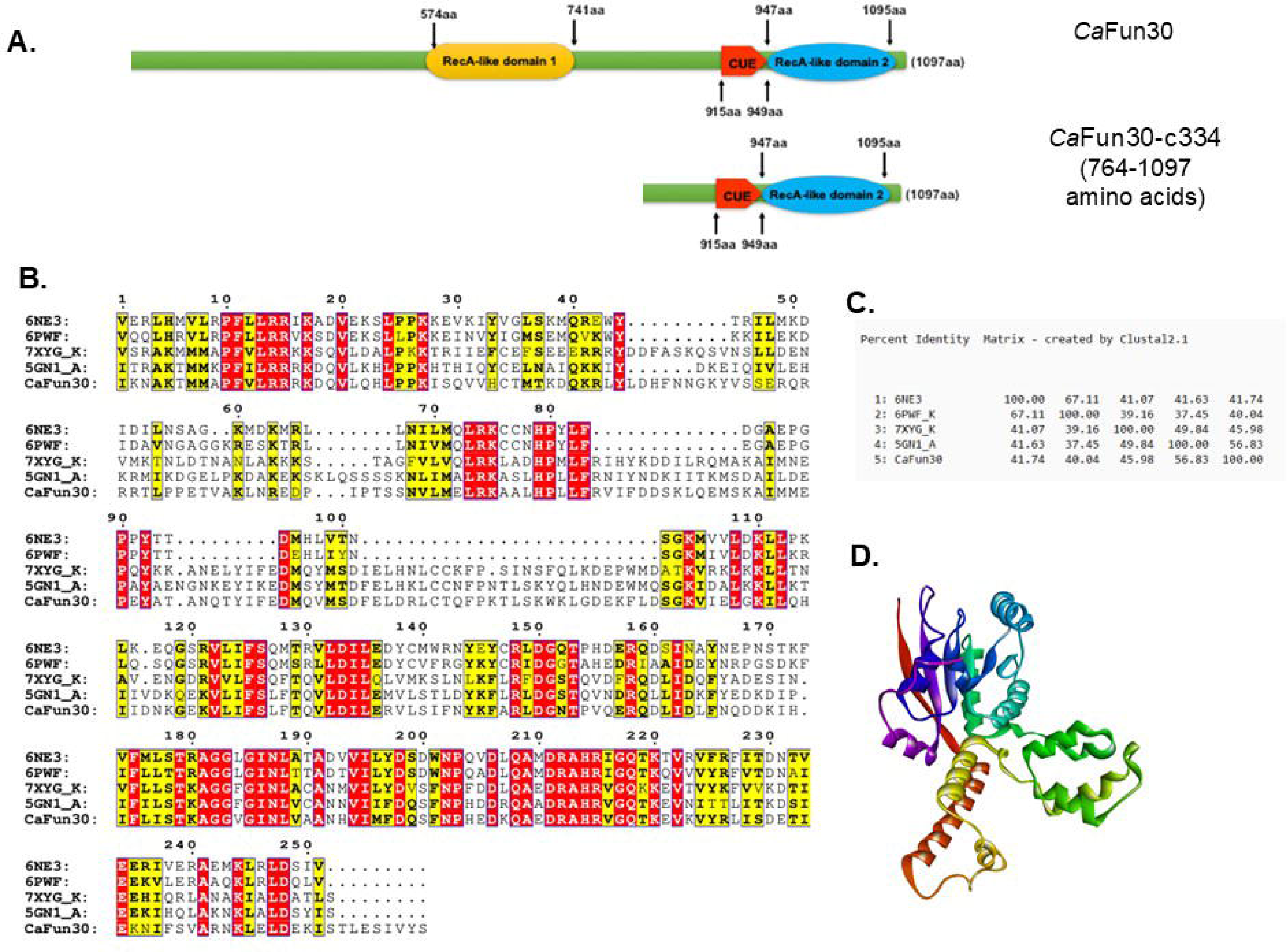
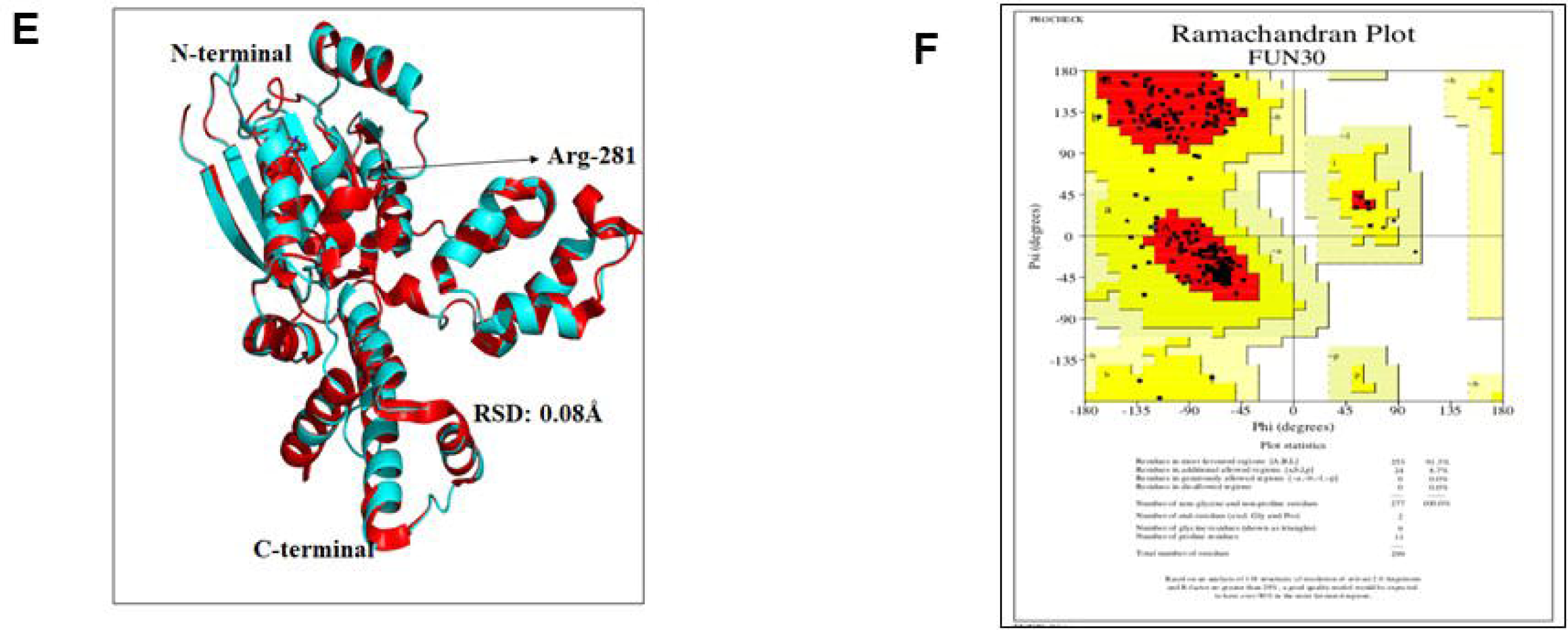
The *Ca*Fun30-c334 structure shares similarities with the *Sc*Fun30 structure. (A). Representation of domains present in *Ca*Fun30 and *Ca*Fun30-c334. *Ca*Fun30 possesses DNA-stimulated ATPase activity as it has both Rec A-like domain 1 and Rec A-like domain 2. *Ca*Fun30-c334 does not show ATPase activity as it possesses only Rec A-like domain 2. (B). Sequence alignment of *Ca*Fun30 with 6NE3 (Snf2h from *H. sapiens*), 6PWF (ISWI from *D. melanogaster*), 7XYG (Fft3 from *D. melanogaster*), and 5GN1 (Fun30 from *S. cerevisiae*). As 5GN1 is a crystal structure of the C-terminal part of Fun30, therefore, only the conserved motifs present in this region are shown. (C). Percent identity between *Ca*Fun30, 6NE3, 6PWF, 7XYG, and 5GN1. (D). Predicted structure of *Ca*Fun30-c334 using homology modelling. (E). The superposition of *Ca*Fun30-c334 and *Sc*Fun30 shows that the structure of the two proteins is similar. (F). Ramachandran plot of the predicted structure of *Ca*Fun30-c334 showing 91.3% residues in the favoured regions.

### Virtual Screening

Life Chemicals Databases [20] with 1,350,000 drug-like and lead-like screening compounds were chosen for virtual screening. Small molecule libraries were prepared by Ligprep of Schrödinger [27,28]. Docking studies were performed using GOLD 5.1 [21], wherein the protein was kept as a rigid molecule, and the number of Genetic Algorithm (GA) runs was set to 10 runs per ligand with the default search algorithm parameters similar to our earlier reports [32,33]. GOLD score, Drug score, and X-SCORE were used to estimate the binding affinity. Interactions between the protein and the compounds were calculated using the Discovery Studio program [30].

### Molecular docking

The selected compounds were further evaluated by molecular docking with AutoDock 4.2 [22]. The complex structure of *Ca*Fun30-c334 and ligands was taken to AutoDock 4.2. Essential hydrogen atoms, atom-type charges, and solvation parameters were added. The whole protein was covered by a cubic grid size (X126 × Y126 × Z126) with a grid spacing of 0.375 Å. The docking simulation was done using the Lamarckian genetic algorithm [34]. Each docking experiment was derived from 40 different conformations with a population size of 300 and 25,000,000 evaluations. The interaction analysis of protein-ligand complexes and their amino acid position with bond distances was calculated and visualised through Discovery Studio [30]. Hydrophobic contacts were analysed using LigPlot+ v. 2.2 [35]. The drug-likeliness of the selected compounds was studied using the SwissADME tool [36] from the Swiss Institute of Bioinformatics (http://www.swissadme.ch/).

### Cloning of CaFUN30 and CaFUN30-c334

The full-length *CaFUN30* gene was amplified from *C. albicans* BWP17 genomic DNA using forward primer 5′-GCTAGCATGAGTTGGTTTAGAAG-3′ and reverse primer 5′-CTCGAGACTATAAACTATTGACTC-3′. The amplified product was cloned between the Nhe1 and Xho1 sites of the pET21c (+) vector. Similarly, the C-terminal fragment *CaFUN30-c334* was amplified from *C. albicans* BWP17 genomic DNA using forward 5’-GGAATTCCATATGGAGAATGATCACAACCCATTG-3’ and reverse 5’-CGCGGATCCTCAACTATAAACTATTGACTCTAATGTTG-3’ primers. The amplified product was cloned into the pCOLD vector.

### Purification of *Ca*Fun30 and *Ca*Fun30-c334

For expression of both proteins, *E. coli* BL21 (DE3) Rosetta cells were transformed with the respective expression plasmid and plated on LB agar supplemented with 100 μg/ml ampicillin and 33 μg/ml chloramphenicol. A single colony was used to inoculate 10 ml LB medium containing the same antibiotics, and the culture was grown overnight at 37°C with shaking at 220 rpm. This primary culture was used to inoculate 1 L LB medium containing 100 μg/ml ampicillin and 33 μg/ml chloramphenicol. The cells were grown at 37°C and 220 rpm until the OD_600nm_ reached 0.6, at which point protein expression was induced with 0.5 mM IPTG. The culture was then incubated at 16°C overnight for protein expression.

For purification of *Ca*Fun30, the cells were harvested after 16 h and resuspended in buffer containing 20 mM Tris-Cl pH 7.5, 350 mM NaCl, 0.1% (v/v) Tween 20, 0.1% (v/v) TritonX-100, 0.1% TritonX-114, 10% glycerol, 0.5 mM PMSF, 5 mM β-mercaptoethanol, 10 mM imidazole, and 0.1 mg/ml lysozyme. After incubation (4°C, 1 h), the cells were sonicated for 5 cycles (15 seconds ON, 45 seconds OFF). The lysate was centrifuged (4°C, 12,000 rpm, 30 min) and supernatant obtained was loaded onto a 1 ml Ni^+2^-NTA column pre-equilibrated with buffer containing 20 mM Tris-Cl pH 7.5, 350 mM NaCl, 0.1% (v/v) Tween 20, 0.1% (v/v) TritonX-100, 0.1% TritonX-114, 10% glycerol, 0.5 mM PMSF, 5 mM β-mercaptoethanol and 10 mM imidazole. The column was washed sequentially with buffer containing 30 mM and 50 mM imidazole. The protein was eluted in 1 ml fractions with buffer containing 100 mM, 200 mM, and 300 mM imidazole. The fractions were analysed on SDS-PAGE, and protein-containing fractions were pooled together.

For purification of *Ca*Fun30-c334, the cells were harvested after 16 h and resuspended in buffer containing 20 mM Tris-Cl pH 8.5, 350 mM NaCl, 0.1% (v/v) Tween 20, 0.1% (v/v) TritonX-100, 0.1% TritonX-114, 10% glycerol, 0.5 mM PMSF, 5 mM β-mercaptoethanol, 10 mM imidazole, and 0.1 mg/ml lysozyme. After incubation (4°C, 1 h), the cells were sonicated for 5 cycles (15 seconds ON, 45 seconds OFF). The lysate was centrifuged (4°C, 12,000 rpm, 30 min) and supernatant obtained was loaded onto a 1 ml Ni^+2^-NTA column pre-equilibrated with buffer containing 20 mM Tris-Cl pH 8.5, 350 mM NaCl, 0.1% (v/v) Tween 20, 0.1% (v/v) TritonX-100, 0.1% TritonX-114, 10% glycerol, 0.5 mM PMSF, 5 mM β-mercaptoethanol, and 10 mM imidazole. The column was washed sequentially with buffer containing 30 mM and 50 mM imidazole. The protein was eluted in 1 ml fractions with buffer containing 100 mM, 200 mM, and 300 mM imidazole. The fractions were analysed on SDS-PAGE, and protein-containing fractions were pooled together.

### Determination of binding affinity of ligands

The binding affinity was determined using biolayer interferometry (BLI) on a Sartorius Octet R8 system with Ni^+2^-NTA sensors.

*Ca*Fun30-c334 and *Ca*Fun30 (100 ng/200 μl) were immobilised on the sensor, and each ligand was titrated at different concentrations to determine the binding affinity. 1X Phosphate-buffered Saline (PBS) pH 7.4 with 2 mM β-mercaptoethanol was used as the running and dissociation buffer. Compound stock solutions were prepared in 100% DMSO, and working dilutions were made in the running buffer. Running buffer alone was used for the control sensor to normalise the data. The association and dissociation kinetics were studied for 120 s each, and data were acquired through Octet BLI Discovery 12.2 software. Data analysis was performed using Octet BLI analysis 12.1 software.

The interaction between the ligand (A), immobilised on the biosensor surface, and the analyte (B), present in solution, was modelled as a simple 1:1 binding reaction:

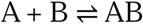

Where complex (AB) formation is characterised by the association rate constant (k_a_) for the forward reaction and the dissociation rate constant (k_d_) for the reverse reaction.

K_D_ is the affinity constant, or equilibrium dissociation constant, which measures how tightly the ligand binds to its analyte. It represents the ratio of the on-rate to the off-rate and can be calculated using k_a_ and k_d_:

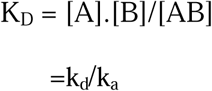

K_D_ is expressed in molar units (M). The K_D_ corresponds to the concentration of analyte at which 50% of ligand binding sites are occupied at equilibrium or the concentration at which the number of ligand molecules with analyte bound equals the number of ligand molecules without analyte bound. The final model fitting equation:

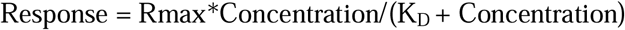

When the highest possible analyte binding is reached, it is considered the maximum binding signal, or Rmax.

### Estimation of ATP hydrolysis

The ATPase activity of *Ca*Fun30 was determined by coupling ATP hydrolysis to NADH oxidation according to the previously standardised protocol [37]. Briefly, 1 μM of protein was incubated with 2 μM fork DNA in the presence of 1× REG buffer containing 25 mM Tris-OAc pH 7.5, 6 mM Mg(OAc)_2_, 60 mM CH_3_COOK, 5 mM β-mercaptoethanol, 1.4 mg/ml phosphoenolpyruvate (PEP) and 10 units each of pyruvate kinase (PK) and lactate dehydrogenase (LDH) in a 250 μl reaction volume. The reactions contained 2 mM ATP and 0.1 mg/ml NADH. The final concentration of the inhibitor was 2 μM. The reaction mixture was incubated at 37°C, and the change in absorbance was recorded at 340 nm as a function of time. The concentration of NADH oxidised was calculated using the molar extinction coefficient of NADH as 6.3 mM^−1,^ wherein 1 mol of ATP hydrolysed results in 1 mol of NADH oxidised.

### Estimation of minimum inhibitory concentration (MIC_50_ and MIC_80_)

For *in vitro* validation of the selected compounds, a cell viability assay was adapted from the previously mentioned protocol [15]. Briefly, the primary culture was grown overnight, and the secondary culture was grown for 6 h at 30°C. For the antifungal assay, two-fold serial dilutions of the stock solution of F1853-0039 were prepared in YEPD medium from 0 to 300 μM (final volume = 100 µl) in a 96-well plate. Log phase cells from the secondary culture were first diluted in 0.9% saline to an OD_600_ nm equal to 0.01. 100 µl of this inoculum was added to each well except for the sterility control, where saline was added to the well. After 16 h incubation at 30°C, OD_600nm_ was monitored using BioTek Cytation 5 imaging reader.

MIC_50_ and MIC_80_ values were calculated by determining the concentration of compounds that lead to an inhibition of cell growth by 50% or 80% as compared with cell growth in the absence of compounds in the growth medium.

To study synergism, the interaction of F1853-0039 with the other antifungal drugs was evaluated using the checkerboard method. The results are represented by the fractional inhibitory concentration index (FICI), with FICI = (MIC of antifungal agent in combination/MIC of antifungal agent alone) + (MIC of F-1853-0039 in combination/MIC of F-1853-0039 alone). FICI ≤ 0.5 shows synergism, 0.5 ≤ FICI ≤ 4.0 shows indifference, and FICI> 4.0 shows antagonism.

To study potential resistance development, an antibiotic susceptibility assay was performed. Cells were subcultured daily in YEPD containing 1 µM F1853-0039 for two weeks under standard growth conditions. Following this period, the minimum inhibitory concentration was determined as described above.

### Toxicity assay on mammalian cells

The susceptibility of human cells towards F1853-0039 was analysed by MTT assay. Human THP-1 cells were seeded at a 6000 cells/well density in a 96-well plate. Phorbol-12-myristate-13-acetate (PMA) (50 ng/ml for 48 h) was added to THP-1 cells to induce their differentiation into macrophage-like cells.

The cells were then treated with different concentrations (0-5000 µM) of F1853-0039 for 24 h. After treatment, MTT was added to a final concentration of 0.5 mg/ml and incubated for 3 h at 37°C for the formation of formazan crystals in living cells. Later, the solution was discarded, and the formazan crystals were dissolved in DMSO (100 µl/well), and absorbance was measured at 570 nm in a BioTek Cytation 5 imaging reader. Cell survivability was checked by measuring the IC_50_ after plotting a nonlinear fit graph of log (inhibitor) vs normalised response in GraphPad Prism version 10.4.1.627.

### Co-culture assay

Co-culture assay was performed as mentioned previously [18]. *C. albicans* cells (100 million) were labelled with 100 μg/ml Calcofluor white (CFW) for 30 min at room temperature, followed by two washes with phosphate-buffered saline at 2000 × g for 5 min each at 4°C. THP-1 cells (0.3 million) were grown on coverslips and incubated with CFW-labelled BWP17 and SN152 strains at M.O.I (1:5) for 1 h at 37°C in a CO_2_ incubator.

Subsequently, 300 μM F1853-0039 was added to the culture and again incubated at 37°C in the CO_2_ incubator. Cells were harvested at 0 h, 2 h, and 6 h of incubation, washed with PBS, and fixed with methanol. The uptake of stained BWP17 and SN152 by THP-1 cells and hyphae formation was examined using a confocal microscope (Nikon Eclipse Ti2 Laser Scanning microscope) under a 60X oil immersion objective.

Quantitation of CFW stain was done using the “intensity line profile” feature of the NIS-Elements AR (Advanced Research) software. Briefly, a circle was drawn around each cell, and the software showed the pixels along this circle. The numbers were individually averaged for all the cells and plotted using GraphPad Prism 10.4.1.627

### Quantitative real-time PCR (qPCR)

A secondary culture of BWP17 and SN152 cells was grown till OD 600 nm reached 0.6. The cells were incubated with 300 µM F1853-0039 for another 6 h. After incubation, the untreated and F1853-0039-treated cells were collected by centrifugation and washed twice with PBS. The cells were lysed in 500 µl TRIzol reagent (Qiagen, Hilden, Germany) by vortexing with glass beads. 200 µl of chloroform was then added, and the solution was centrifuged at 12,000 rpm for 15 min. RNA was precipitated by adding 1 mL of ethanol to the supernatant. The integrity and quantity of the isolated RNA were estimated using a NanoDrop Spectrophotometer (Thermo Fisher Scientific, USA). cDNA was prepared using 1 µg of total RNA according to the manufacturer’s protocol using the cDNA synthesis kit from Thermo Fisher Scientific, USA. SYBR Green (ABI) was used for the quantitative assessment of the mRNA level. 1 μl of diluted cDNA was used for these assays. The levels were quantitated using gene-specific primers (Supplementary Table 2), and 18S ribosomal RNA (*RDN18*) was taken as an internal control. The PCR parameters using real-time ABI 7500 Fast Cycler are provided in Supplementary Table 3. The fold change was calculated using the delta-delta Ct method [38] where; ΔCt sample = Ct (gene of interest) – Ct (internal control), ΔCt non-target control = Ct (gene of interest) – Ct (internal control), ΔΔCt = ΔCt sample - ΔCt non-target control and fold expression = 2^-ΔΔCt^.

### Statistical analysis

All qPCR and ChIP experiments are reported as the average ± standard error of mean (SEM) of three independent (biological) experiments unless otherwise specified. Each independent experiment was performed with at least two technical replicates. The statistical significance (p-value) was calculated using a paired t-test available in GraphPad Prism version 10.4.1.627. The differences were considered significant at p < 0.05.

## RESULTS

### The *Ca*Fun30 structure shares similarities with the *Sc*Fun30 structure

Understanding the three-dimensional structure of *Ca*Fun30 provides greater insight into its function. However, the X-ray structure of the *Ca*Fun30 protein is not available in the protein data bank, and therefore, computational modelling was used for predicting the 3D structural model. The crystal structure of the Fun30 ATPase domain from *S. cerevisiae* S288C, solved at 1.95Å resolution (PDB: 5GN1)[25] shares a sequence identity of 55 % (Fig.1A-C).

Sequence analysis using MUSCLE showed that this elucidated structure contains only motifs IV-VI (Fig. 1B) and is referred to as *Ca*Fun30-c334 throughout the text. This region of the protein lacks ATPase activity. Modeller software was used to model the 3D structural model of *Ca*Fun30-c334 using 5GN1 as the structure template. The structure was prepared using the Protein Preparation Wizard of Schrodinger software to remove bad contacts (Fig. 1D).

The *Ca*Fun30-c334 model prepared by Structural Superposition of Fun30 structure (PDB: 5GN1) for 31-239 residues revealed RMSD of 0.08 Å (Fig. 1E). The quality of the model was evaluated using PROCHECK to confirm the stereochemical quality of the modelled structure. Ramachandran plot analysis revealed that 100% of residues of *Ca*Fun30-c334 (Fig.1F) were found in the favoured regions, ensuring that the structure is reliable for further studies. The Arg-281 residue in the arginine finger in motif IV is conserved in the Fun30 ATPase domain (PDB: 5GN1) (Fig. 1E). Therefore, the active site was defined as 10Å around the Arg-281 residue.

### Ten ligands were identified as potential inhibitors of *Ca*Fun30-c334

GOLD 5.1 program was used for virtual screening of the Life Chemicals compound library. The number of Genetic Algorithm (GA) runs was set to 10, with a population size of 100, and the number of operations was set to 100,000, with selection pressure maintained at 1.1. The default GoldScore scoring function was employed to screen the library of compounds from the Life Chemicals database [20]. Library screening was done to filter the non-docked compounds. After filtering, the remaining compounds were used for detailed docking. GOLD 5.1 program [21] uses a genetic algorithm for docking ligands into protein binding sites to explore the full range of ligand conformational flexibility with partial flexibility of protein. The docking procedure consists of three interrelated components: (i) identification of the binding site, (ii) a search algorithm to effectively sample the search space (the set of possible ligand positions and conformations on the protein surface), and (iii) a scoring function. The best 10 ligands were chosen based on the highest scores obtained from the genetic docking algorithm GOLD and the consensus scoring program X-Score [39] (Table 1).

**Table 1:** Docking of *Ca*Fun30-c334 with the identified top 10 inhibitors and their estimated binding affinities using the GOLD 5.1 [21] program.

| # | Ligands | GOLD score | XSCORE<br>(kcal/mol) | -log(K<sub>d</sub>) | Drugscore<br>(kcal/mol) |
| --- | --- | --- | --- | --- | --- |
| 1. | F1853-0039 | 76.27 | -7.60 | 5.58 | -83 |
| 2. | F1067-0218 | 79.1 | -8.23 | 6.03 | -89 |
| 3. | F2716-0594 | 73.4 | -7.65 | 5.88 | -76 |
| 4. | F3385-2833 | 78.0 | -8.95 | 6.56 | -82 |
| 5. | F3139-2622 | 78.3 | -8.15 | 5.98 | -84 |
| 6. | F2814-1613 | 75.06 | -7.68 | 5.90 | -82 |
| 7. | F5670-0144 | 71.6 | -7.45 | 5.46 | -73 |
| 8. | F0196-0182 | 80.4 | -8.02 | 5.88 | -88 |
| 9. | F0196-0638 | 77.4 | -8.05 | 5.90 | -85 |
| 10. | F3407-0022 | 72.7 | -7.76 | 5.69 | -74 |

Molecular docking was further confirmed using AutoDock 4.2 with 40 GA runs, a population size of 300 and 25,000,000 energy evaluations per run. The mutation and crossover rates were set at 0.02 and 0.80, respectively. The grid spacing was maintained at 0.375 Å, and the protein was treated as rigid while ligand torsional flexibility was allowed.

The ligand and protein structures were first converted to AutoDock 4.2-compatible input formats (map files and pdbqt) using AutoDock 4.2 Tools. Docking was performed with the Lamarckian Genetic Algorithm [34] to determine the optimal docking position between the ligands and the macromolecules. The results were then evaluated with an empirical binding free energy function and plotted in LigPlot+ (Table 2; Fig. 2). The data generated showed that the range of X-Score for the top 10 compounds from the Life Chemicals library ranged between -7.45 kcal/mol to -8.95 kcal/mol, and their binding affinity calculated in AutoDock 4.2 ranged between -15.29 kcal/mol to -5.72 kcal/mol. Of these 10, only 7 were procurable.

**Figure 2.**
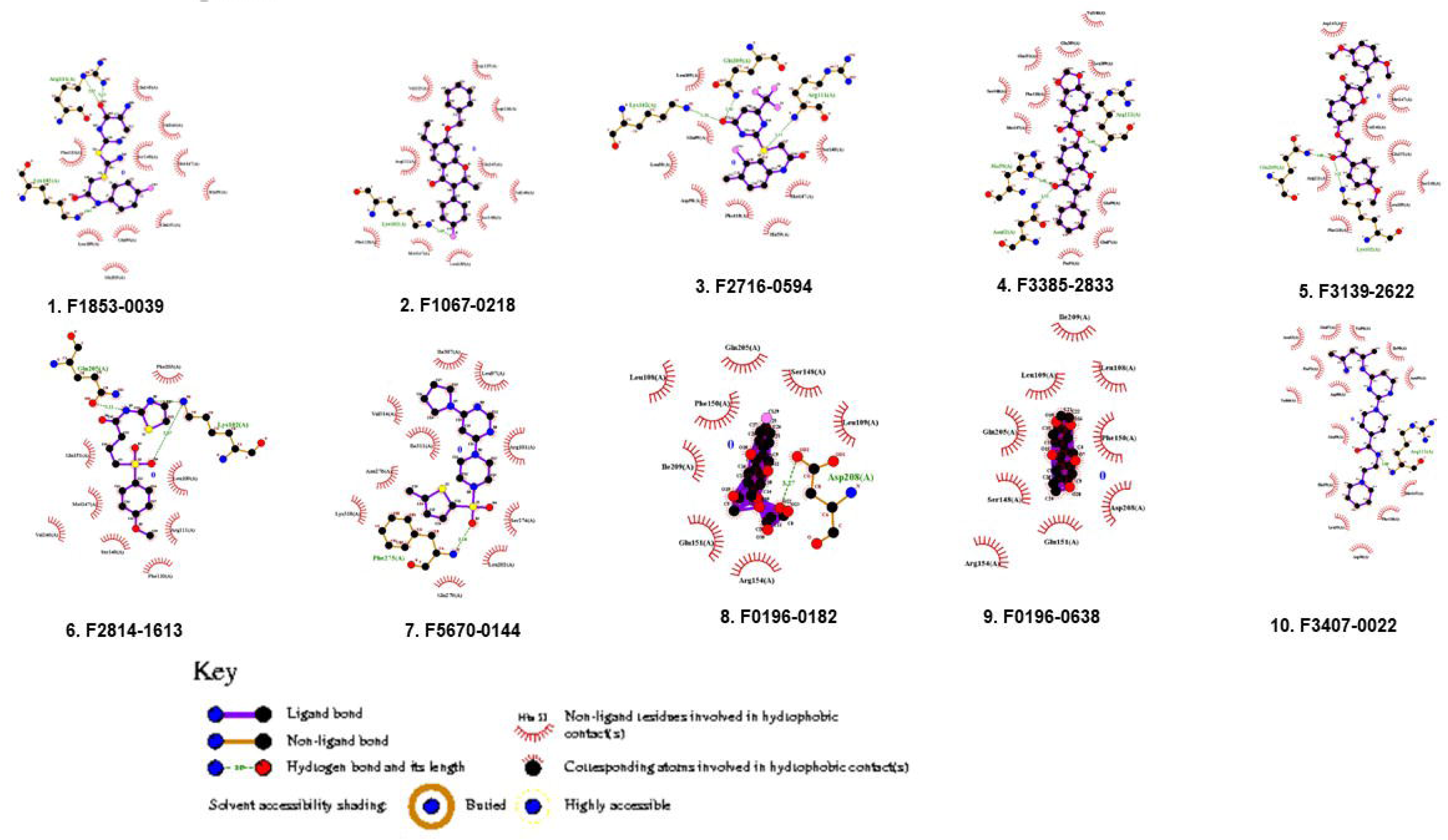
Ten ligands were identified as potential inhibitors of *Ca*Fun30-c334. Screening of the Life Chemicals [20] database identified ten ligands as potential inhibitors of *Ca*Fun30-c334.

**Table 2:** Docking of *Ca*Fun30-c334 with the identified top 10 inhibitors and their estimated binding affinities using Autodock 4.2. [22].

| # | Ligands | Binding Energy (kcal/mol) | Ligand Efficiency | Inhibition Constant ($K_i$ or $K_d$) μM | Inter-molecular Energy | VE | Electrostatic energy | Total internal energy | Torsional energy | Un-bound energy |
| --- | --- | --- | --- | --- | --- | --- | --- | --- | --- | --- |
| 1. | F1853-0039 | -6.59 | -0.29 | 14.7 μM | -8.98 | -7.79 | -1.19 | -1.53 | -2.39 | -1.53 |
| 2. | F1067-0218 | -6.67 | -0.23 | 12.81 μM | -8.17 | -7.92 | -0.25 | -1.5 | 1.49 | -1.5 |
| 3. | F2716-0594 | -5.72 | -0.24 | 64.19 μM | -7.21 | -7.17 | -0.05 | -1.56 | 1.49 | -1.56 |
| 4. | F3385-2833 | -7.99 | -0.28 | 1.38 μM | -9.19 | -8.96 | -0.23 | -0.93 | 1.19 | -0.93 |
| 5. | F3139-2622 | -6.9 | -0.21 | 8.75 μM | -9.29 | -8.91 | -0.38 | -1.0 | 2.39 | -1.0 |
| 6. | F2814-1613 | -6.05 | -0.27 | 36.96 μM | -8.13 | -7.89 | -0.24 | -0.56 | 2.09 | -0.56 |
| 7. | F5670-0144 | -6.76 | -0.26 | 11.05 μM | -7.96 | -7.96 | 0.01 | -0.93 | 1.19 | -0.93 |
| 8. | F0196-0182 | -13.57 | -0.45 | 112.78 pM | -17.15 | -17.15 | 0.0 | -1.21 | 3.58 | -1.21 |
| 9. | F0196-0638 | -15.29 | -0.64 | 6.21 pM | -17.38 | -17.38 | 0.0 | 0.0 | 2.09 | 0.0 |
| 10. | F3407-0022 | -7.07 | -0.24 | 6.59 μM | -9.75 | -7.39 | -2.36 | -0.56 | 2.68 | -0.56 |

These seven procurable compounds (F1853-0039, F1067-0218, F2716-0594, F3385-2833, F3139-2622, F2814-1613, F5670-0144) were used for further analysis.

### *In silico* ADME studies show that the ligands of *Ca*Fun30-c334 were potent molecules

The bioavailability radar chart was used to investigate the physicochemical and drug-likeness properties of a molecule through the prediction of six physicochemical descriptors: lipophilicity (LIPO, the partition coefficient between n-octanol and water log Po/w value should be between −0.7 and +5.0), size (the acceptable molecular weight lies between 150 and 500 Da), polarity (POLAR), topological polar surface area (TPSA between 20 and 130 Å2), insolubility (INSOL, the decimal logarithm of the molar solubility in water log S should not exceed 6), unsaturation (UNSAT, fraction of carbons in the sp3 hybridization ≥0.25), and flexibility (FLEX, the number of rotatable bonds should not be greater than 9). The optimal physicochemical range on each axis is depicted as a pink area in which the radar plot of the molecule must fall entirely to be considered drug-like (Supplementary Fig. 1).

The screened results of ADMET are summarised in Supplementary Table 1, revealing various descriptors categorised under the above-mentioned properties. The physicochemical properties of all the selected compounds were within the given range. It was further observed that all the molecules (except F1853-0039) could be passively absorbed by the gastrointestinal tract and would not cross the blood-brain barrier (Fig. 3). The compounds had the same bioavailability score of 0.55 and did not violate Lipinski’s rule. F2716-0594, F3385-2833, F3139-2622, F2814-1613, and F5670-0144 were positive for drug-likeness in all five different rule-based filters, while F1853-0039 was positive in three, and F1067-0218 was positive in two of the five filters. F1853-0039 was shown to be relatively hydrophilic (TPSA 142.79 Å², Consensus LogP 2.42), whereas the other compounds were predominantly hydrophobic (Consensus LogP > 3.5). This difference is important because hydrophilicity can enhance aqueous solubility and influence distribution and clearance, while hydrophobicity is often associated with better membrane permeability but poorer solubility.

**Figure 3.**
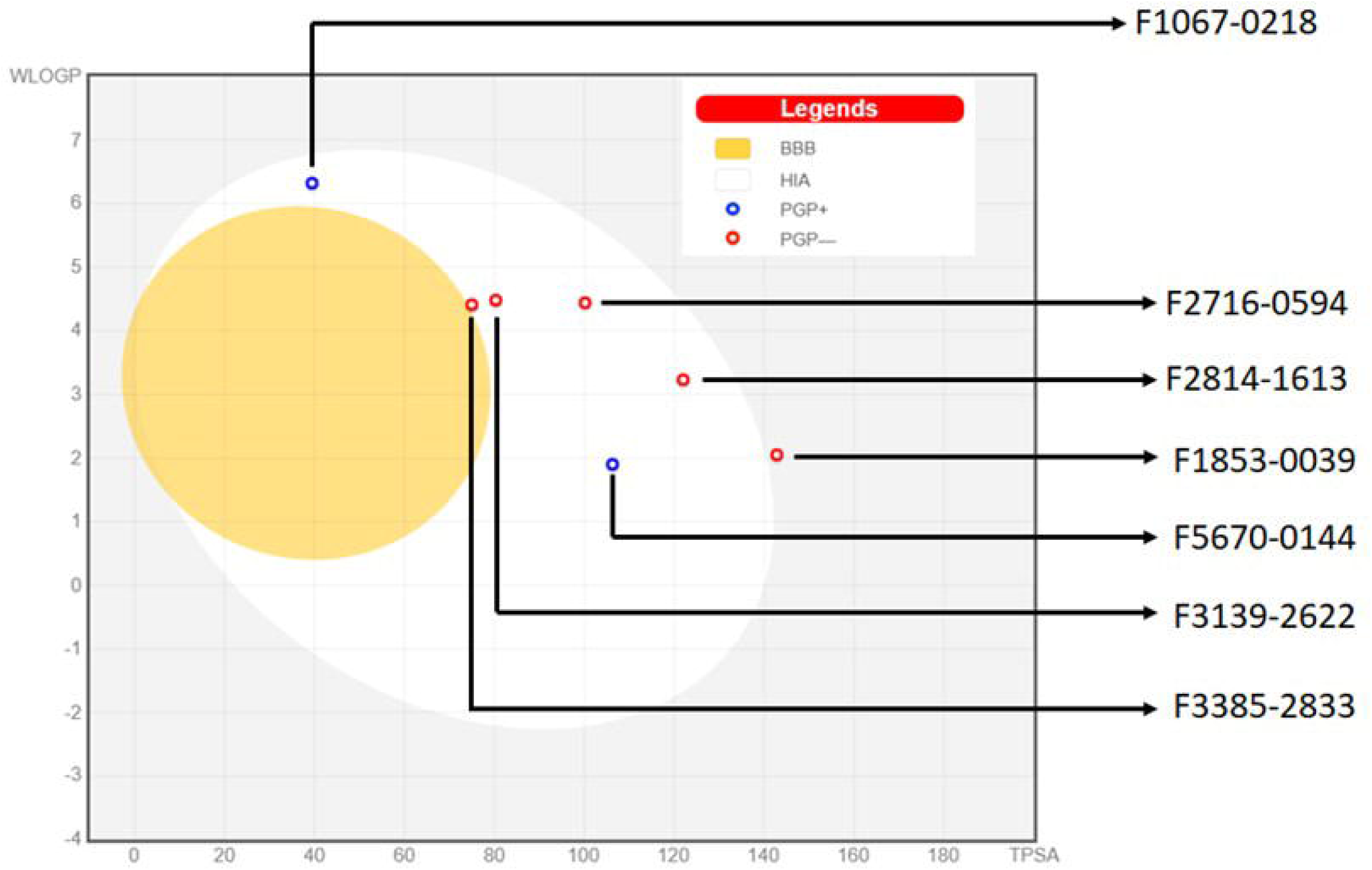
*In silico* ADME studies show that the ligands of *Ca*Fun30-c334 were potent molecules. Boiled egg plot for the ten potential inhibitors of *Ca*Fun30-c334. None of the molecules are predicted to penetrate the brain as they lie outside the yellow (yolk) region. F1853-0039 is predicted not to be absorbed as it lies outside the white region. F5670-0144 and F1067-0218 are predicted to be actively effluxed by P-glycoprotein.

Therefore, the balance of these properties is critical for optimising pharmacokinetic behaviour. Thus, the ADME analysis suggested that the selected compounds were potent molecules and could be effective drugs.

### F1853-0039 shows the highest affinity for *Ca*Fun30-c334 and *Ca*Fun30

Of the seven compounds procured for analysis, only 5 (F1853-0039, F1067-0218, F2716-0594, F3385-2833, and F3139-2622) were soluble in DMSO. The interaction of these compounds with purified *Ca*Fun30-c334 and *Ca*Fun30 was studied using BLI. The compounds F1853-0039, F1067-0218, and F2716-0594 showed significant binding for both *Ca*Fun30-c334 (Fig. 4A-C) and *Ca*Fun30 (Fig. 4D-F) proteins, while F3385-2833, F3139-2622, F2814-1613 and F5670-0144 did not show any binding. The dissociation constants (K_d_) for F1853-0039, F1067-0218, and F2716-0594 were calculated to be 870 nM, 18 µM, and 25.1 µM for *Ca*Fun30-c334; and 600 nM, 125 µM, and 680 µM for *Ca*Fun30, respectively. As, F1853-0039 was found to have the highest binding affinity for both *Ca*Fun30-c334 as well as *Ca*Fun30, therefore, this compound was used in further studies.

**Figure 4.**
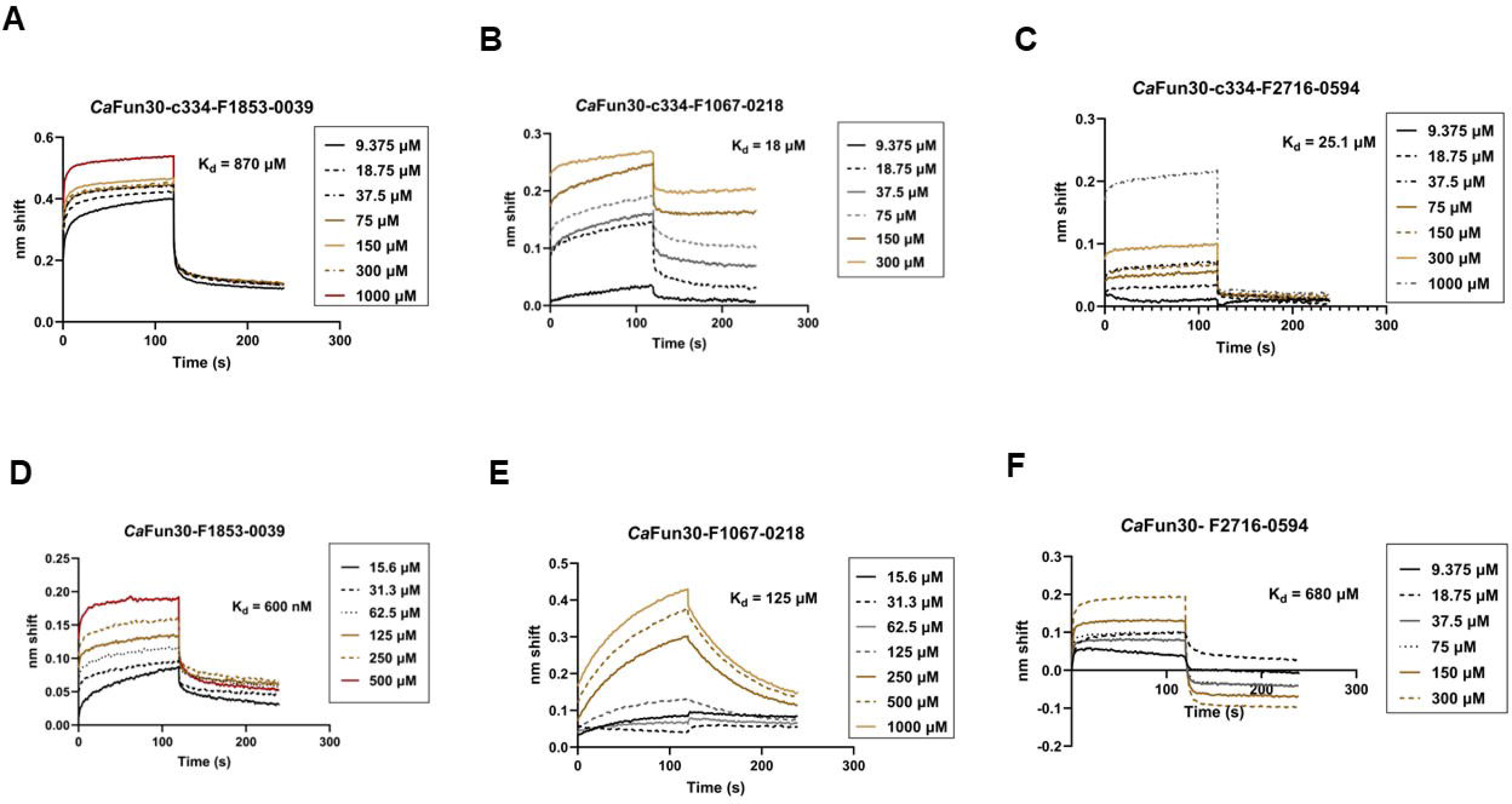
F1853-0039 shows the highest affinity for *Ca*Fun30-c334 and *Ca*Fun30. Binding affinities were estimated for the interaction of (A). F1853-0039; (B) F1067-0218;(C) F2716-0594 with *Ca*Fun30-c334. Binding affinities were estimated for the interaction of F1853-0039; (E) F1067-0218; (F) F2716-0594 with *Ca*Fun30. The binding studies were performed two times, and the best graphs are represented in this figure. 100 ng/200 μl protein was used in this study.

### F1853-0039 inhibits the ATPase activity of *Ca*Fun30 *in vitro*

Fig. 5A provides the structure of F1853-0039. Because Fun30 possesses DNA-stimulated ATPase activity, we next asked whether F1853-0039 can inhibit the ATPase activity of the protein. NADH-coupled ATPase assay showed that F1853-0039 could indeed inhibit the ATPase activity of the protein (Fig. 5B).

**Figure 5.**
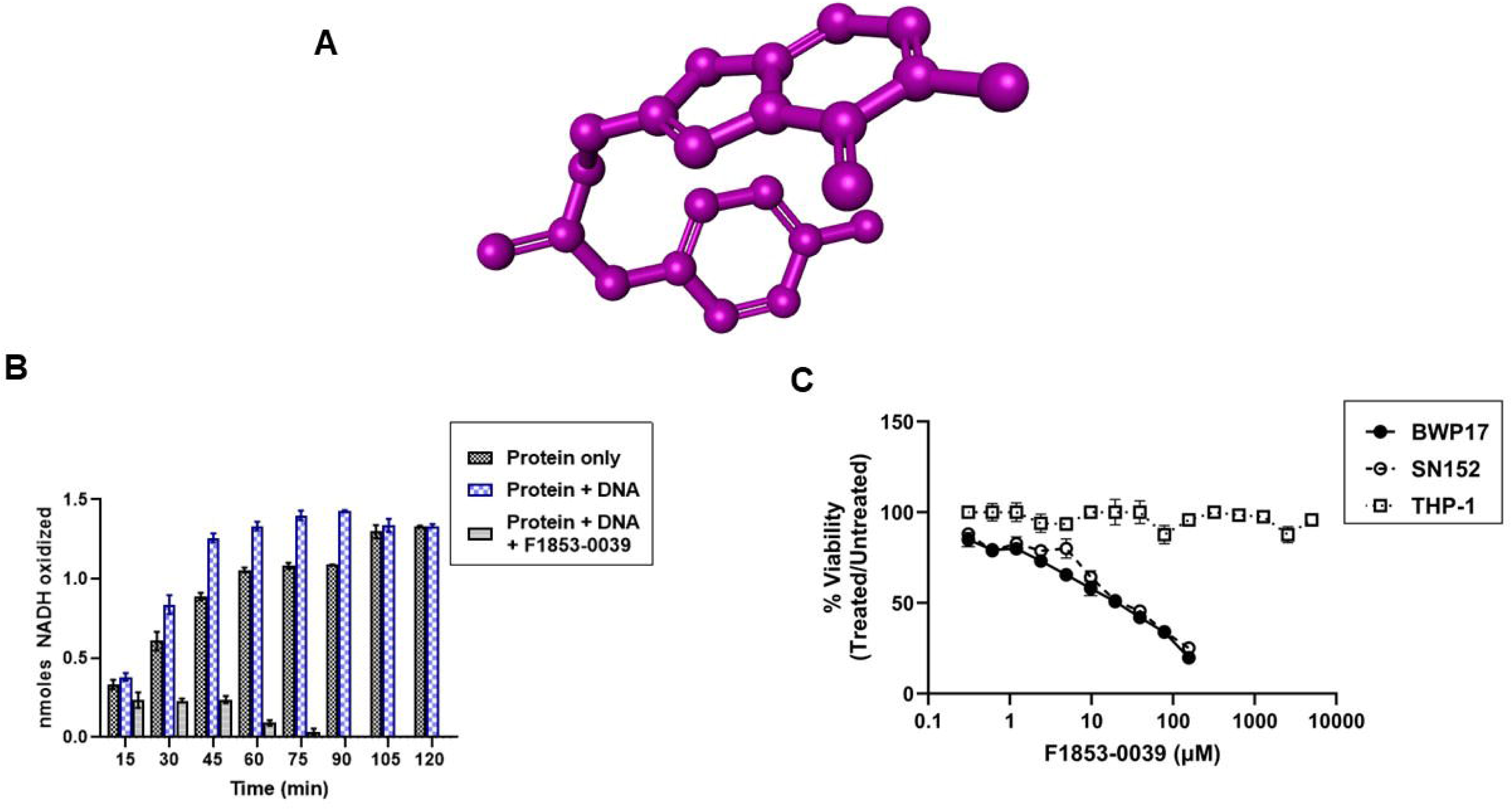
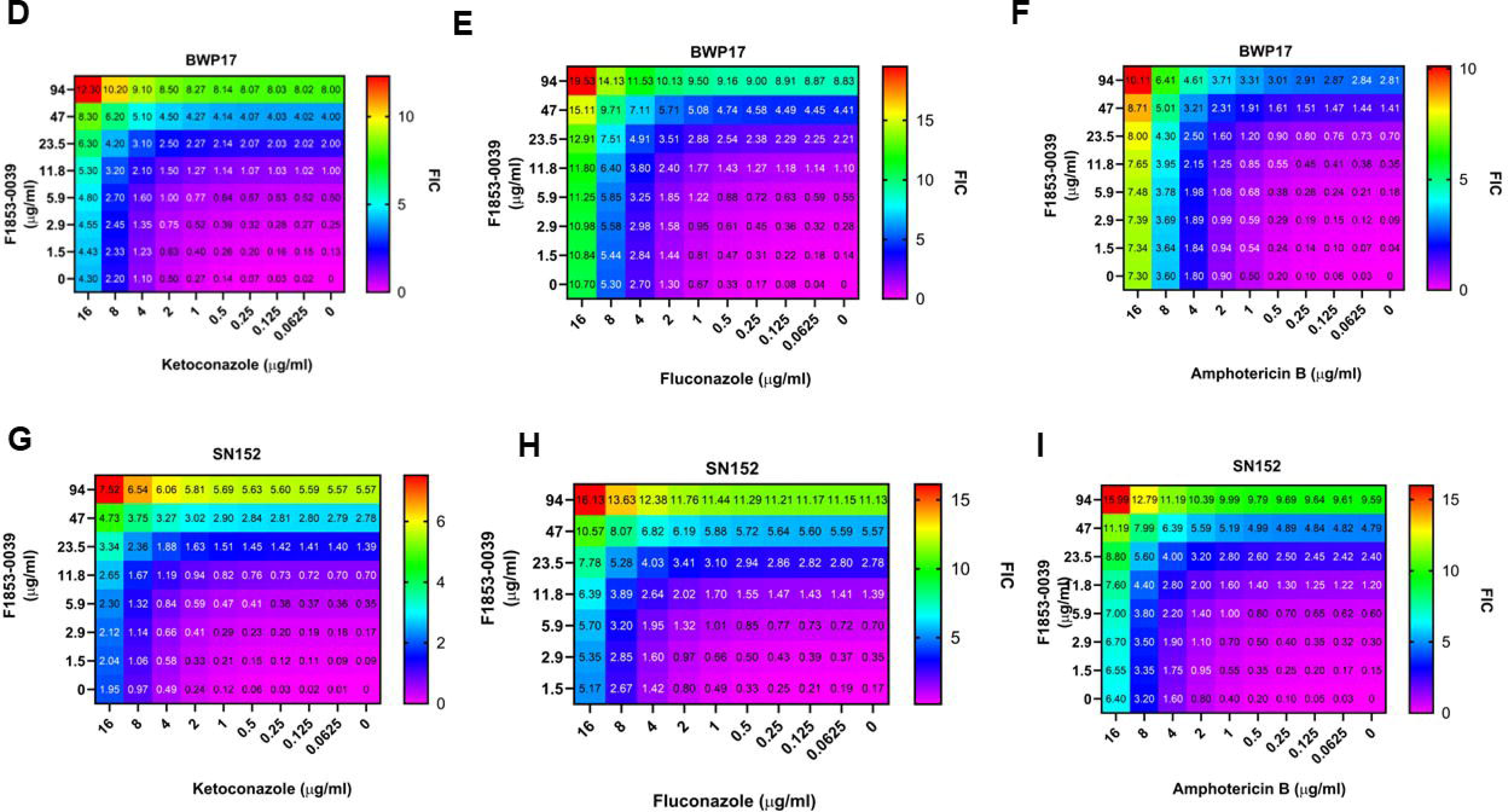
F1853-0039 inhibits the ATPase activity as well as the growth of *Ca*Fun30 *in vitro*. (A). Structure of F1853-0039. (B). The ATPase activity of purified *Ca*Fun30 was estimated in the absence and presence of 2 μM F1853-0039 using the NADH oxidation assay. In this assay, 1 mole of ATP hydrolysed is equal to 1 mole of NADH oxidised. The data is presented as an average ± SD of two independent experiments. (C). The effect of F1853-0039 on BWP17, SNF152, and human cell line THP-1 was monitored by incubating an increasing concentration of the inhibitor as explained in Materials and Methods. The cell viability of *C. albicans* strains was monitored by measuring absorbance at 600 nm. MTT assay was used to measure the cell viability of THP-1 cells. The data is presented as the average ± SEM of three independent replicates. The effect of F1853-0039 in combination with (D) ketoconazole, (E) fluconazole, and (F) amphotericin B was studied in BWP17 cells. The effect of F1853-0039 in combination with (G) ketoconazole, (H) fluconazole, and (I) amphotericin B was studied in SN152 cells. FICI was calculated as explained in the materials and methods. Synergism <0.5; antagonism>4; additive or no effect 0.5-4.

### F1853-0039 inhibits *C. albicans* growth *in vitro*

To understand the cytotoxicity of F1853-0039, a cell viability assay was performed using BWP17 and SN152 strains of *C. albicans*. The cells were treated with increasing concentrations of F1853-0039 for 16 h at 30 °C, and cell viability was estimated by measuring the optical density at 600 nm. For BWP17, the MIC_50_ value obtained for F1853-0039 was 15.8 µM (5.8 μg/ml) and the MIC_80_ value was 25.3 μM (9.3 μg/ml) (Fig. 5C). For SNI52, the MIC_50_ value obtained for F1853-0039 was 19.1 µM (7.0 μg/ml) and MIC_80_ value was 30.5 μM (11.2 μg/ml) (Fig. 5C). Thus, the MIC values of F1853-0039 was almost similar in BWP17 and SN152, suggesting its efficiency was same for different *C. albicans* strains.

### F1853-0039 does not affect the growth of THP-1 cells

On infection, *Candida albicans* reside within the macrophages. Hence, the effect of F1853-0039 on the human cell line-derived macrophages was assessed using THP-1 cells, which are monocytes isolated from the peripheral blood of a human leukaemia patient. The cells were treated with PMA to induce differentiation into macrophages. Subsequently, the cells were incubated with increasing concentrations (0-5000μM) of F1853-0039 for 24 h. The cell viability was measured using the MTT assay. We found that F1853-0039 did not affect the viability of the host cells (Fig. 5C).

Thus, F1853-0039 appears to be more effective against *C. albicans* as compared to the mammalian cell line.

### F1853-0039 shows synergism with azoles and amphotericin B

To understand synergism with existing antifungal drugs, the effect of the combination of amphotericin, fluconazole, and ketoconazole with F1853-0039 was evaluated via the checkerboard method. Based on checkerboard association testing, amphotericin, fluconazole, and ketoconazole exhibited a synergistic activity against BWP17 and SN152 (Fig. 5D-I). It was noted that ketoconazole MIC decreased from 3.7 µg/ml to 0.5 µg/ml (for BWP17) (Fig. 5D), with FICI of 0.26, and from 8.2 µg/ml to 1 µg/ml (for SN152), with FICI of 0.2 when combined with 1.5 µg/ml F1853-0039 (Fig. 5E). The MIC of fluconazole decreased from 1.5 µg/ml to 0.125 µg/ml (for BWP17) (Fig. 5F), with FICI of 0.22, and from 3.2 µg/ml to 0.125 µg/ml (for SN152), with FICI of 0.21 when combined with 1.5 µg/ml F1853-0039 (Fig. 5G). And amphotericin MIC decreased from 2.2 µg/ml to 0.5 µg/ml (for BWP17) (Fig. 5H), with FICI of 0.24, and from 2 µg/ml to 0.25 µg/ml (for SN152), with FICI of 0.25 when combined with 1.5 µg/ml F1853-0039 (Fig. 5I).

### F1853-0039 inhibits the formation of hyphae in co-culture

To understand whether F1853-0039 inhibits hyphae formation, cells were grown in RPMI media supplemented with FBS. Examination of cells showed that hyphae formation was reduced in both BWP17 and SN152 cells in the presence of the inhibitor (Supplementary Fig. 3A).

Next, the effect of F1853-0039 on the virulence of *C. albicans* using THP-1 cells was studied. The cells were infected with BWP17 or SN152 (M.O.I = 1:5) and allowed to grow for 1 h. Subsequently, the cells were treated with 300 μM compound (10x MIC_80_) and harvested at 0 h, 2 h, and 6 h time points to observe the *C. albicans* cells and hyphae formation compared to the untreated infected THP-1 cells. The co-culture experiment was performed with 300 µM of the inhibitor molecule, keeping in mind that the concentration used should be more than MIC_80_ to show inhibition of *Candida* growth and hyphae formation, along with exerting minimal effect on THP-1 cells.

We found that the number of *C. albicans* cells per frame and the number of hyphae decreased in the presence of F1853-0039 in both BWP17 and SN152 strains (Fig. 6A-D; Fig. 7A-D). Further, the fluorescence intensity of CFW was found to decrease in both BWP17 and SN152 when treated with F1853-0039 as compared to the untreated condition, suggesting chitin formation is affected by the inhibitor molecule (Fig. 6E; Fig. 7E).

**Figure 6.**
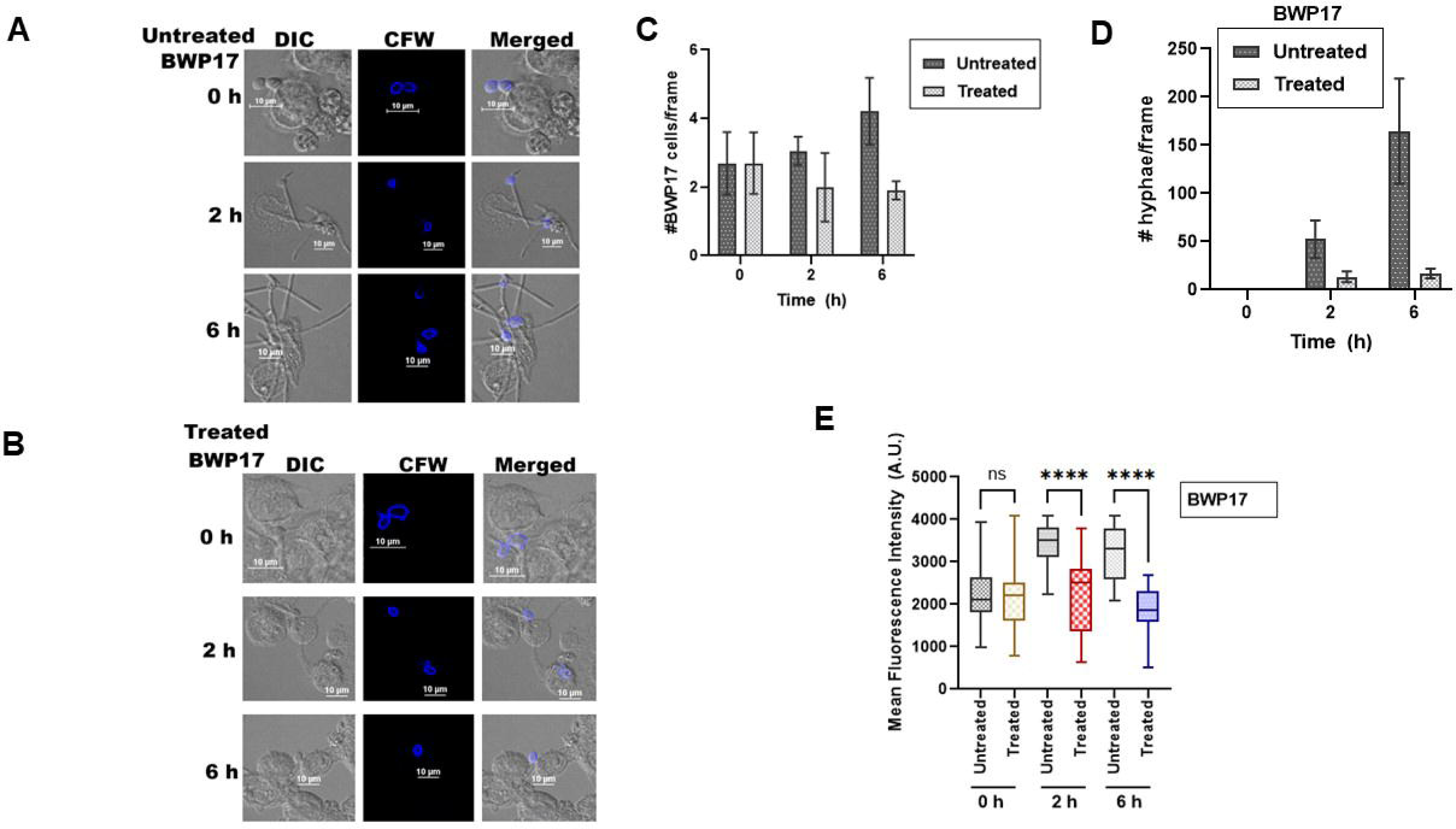
F1853-0039 inhibits the formation of hyphae of BWP17. (A). Growth of BWP17 cells co-cultured with THP-1 cells was monitored at 0 h, 2h, and 6 h in the absence of F1853-0039. (B). Growth of BWP17 cells co-cultured with THP-1 cells was monitored at 0 h, 2 h, and 6 h after treatment with 300 μM F1853-0039. (C). The number of BWP17 cells (as observed by CFW stain) was counted per frame in untreated and treated conditions at 0 h, 2 h, and 6 h after treatment with F1853-0039. (D). The number of hyphae was counted per frame in untreated and treated conditions at 0 h, 2 h, and 6 h after treatment with F1853-0039. (D). The fluorescence intensity of CFW was estimated using the “intensity line profile” feature of the NIS-Elements AR (Advanced Research) software. Briefly, a circle was around each cell, and the pixels along this circle were shown by the software. The numbers were individually averaged for all the cells and plotted using GraphPad Prism version 10.4.1.627. The experiments were done as biological replicates, and the numbers represent the total number of cells calculated from both experiments.

**Figure 7.**
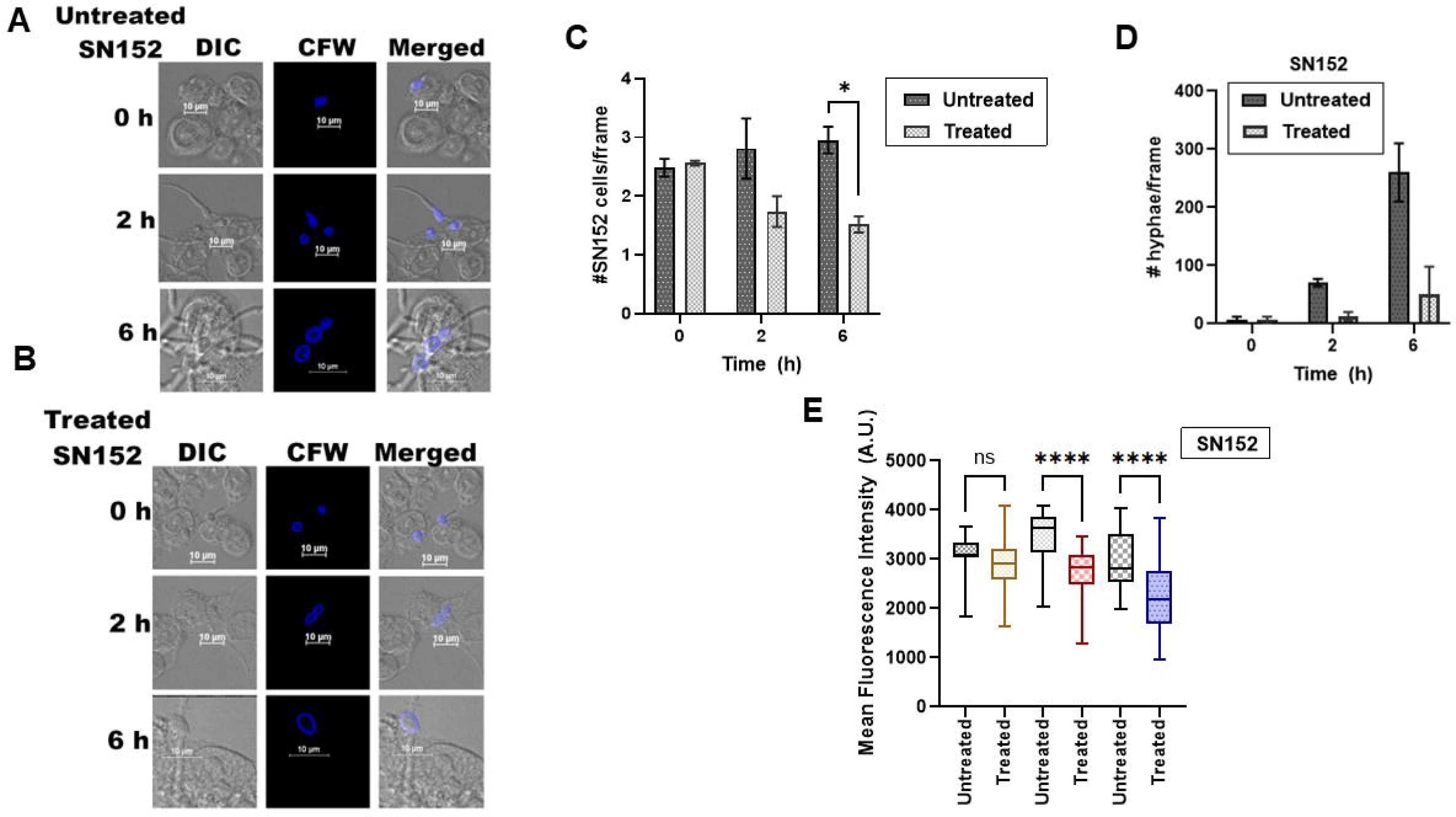
F1853-0039 inhibits the formation of hyphae of SN152. (A). Growth of SN152 cells co-cultured with THP-1 cells was monitored at 0 h, 2 h, and 6 h in the absence of F1853-0039. (B). Growth of SN152 cells co-cultured with THP-1 cells was monitored at 0 h, 2 h, and 6 h after treatment with 300 μM F1853-0039. (C). The number of SN152 cells (as observed by CFW stain) was counted per frame in untreated and treated conditions at 0 h, 2 h, and 6 h after treatment with F1853-0039. (D). The number of hyphae was counted per frame in untreated and treated conditions at 0 h, 2 h, and 6 h after treatment with F1853-0039. (E). The fluorescence intensity of CFW was estimated using the “intensity line profile” feature of the NIS-Elements AR (Advanced Research) software. Briefly, a circle was around each cell, and the pixels along this circle were shown by the software. The numbers were individually averaged for all the cells and plotted using GraphPad Prism.

### Gene expression is altered in the presence of F1853-0039

Previous studies have shown that Fun30 and Rtt109 regulate each other’s transcription. Therefore, we postulated that the expression of *FUN30* and *RTT109* should be downregulated in the presence of F1853-0039. Further, H3K56ac catalysed by Rtt109 is required for *GPI15* expression. This gene encodes for Gpi15, a subunit of the GPI-GnT complex. We hypothesised that the expression of *GPI15* should also be downregulated in the presence of F1853-0039. Hence, the expressions of *FUN30*, *RTT109*, and *GPI15* in F1853-0039-treated BWP17 and SN152 cells were analysed using qPCR. CFW staining indicated alterations in chitin expression, so transcript levels of *CHS1* and *CHS2* were also determined using qPCR. The transcript levels of *FUN30*, *RTT109*, *GPI15*, *CHS1* (P < 0.0001), and *CHS2* (P< 0.0001) were found to be significantly downregulated in F1853-0039 treated cells as compared to the untreated samples, thus, validating our hypothesis (Fig. 8A and B). However, it needs to be noted that *CHS1* and *CHS2* downregulation is only between 40-70% and might not have a dramatic effect on chitin synthesis. Therefore, the reduced chitin staining observed in the presence of the inhibitor could be an interplay of different factors.

**Figure 8:**
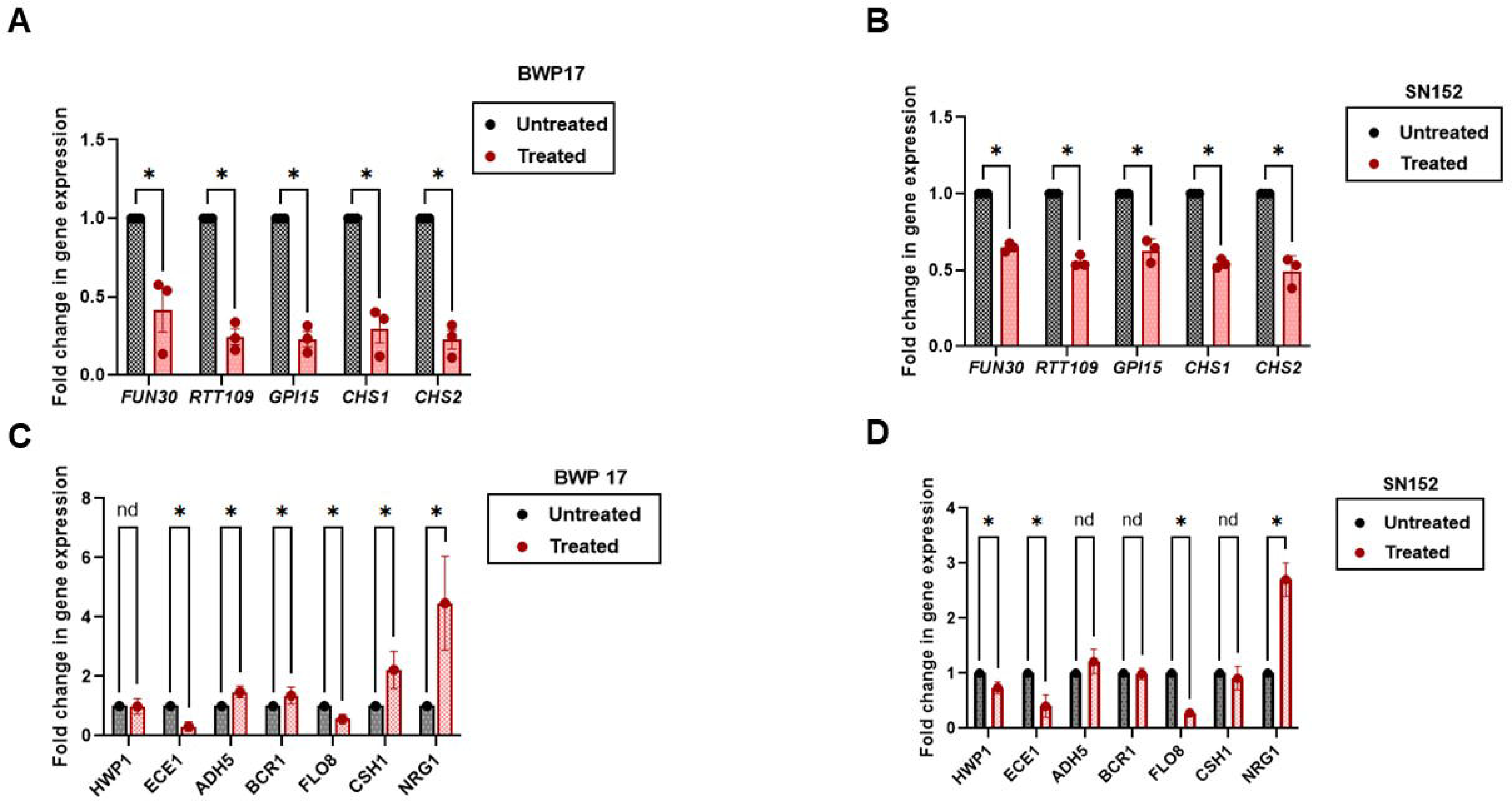
Gene expression is altered in the presence of F1853-0039. Transcript levels of *FUN30*, *RTT109*, *GPI15*, *CHS1*, and *CHS2* were compared between untreated and F185-0039 treated (A). BWP17 cells. (B). SN152 cells. The data is plotted as average ± SEM of three biological replicates. Transcript levels of *ECE1*, *HWP1*, and *NRG1* were compared between untreated and treated (C) BWP17 cells. (D). SN152 cells. The data is plotted as average ± SD of two biological replicates. Star represents p <0.05.

To further confirm the inhibitory effect of F1853-0039 on hyphal formation, we analysed the expression of key hyphae-specific genes in BWP17 and SN152 strains.

Treatment with F1853-0039 resulted in a marked suppression of *ECE1* and *FLO8*, which are typically upregulated during hyphal induction (Fig. 8C and D). Conversely, *NRG1*, a known negative regulator of hyphal development, was significantly overexpressed. Additionally, genes such as *HWP1*, *ADH5*, *BCR1*, and *CSH1* exhibited differential regulation in response to F1853-0039 treatment, indicating transcriptional reprogramming associated with impaired hyphal growth (Fig. 8C and D). The regulation of hyphae-related genes by F1853-0039 further emphasises its potential as a novel antifungal agent targeting pathogenicity pathways.

### Effect of F183-0039 on drug-resistant strains of *C. albicans* and *C. auris* strains

The effect of F1853-0039 on three matched pairs of clinical isolates was also studied using GU4 (drug-sensitive) and GU5 (drug-resistant); DSY 347 (drug-sensitive) and DSY289 (drug-resistant); DSY 544 (Drug-sensitive) and DSY 775 (drug-resistant) [40]. We found that only GU5 was not responsive to F1853-0039 (Supplementary Fig. 2A-C). We also tested the inhibitor on two drug-resistant strains of *C. auris* (NCCPF 470156 and NCCPF 470157) [41]. NCCPF 470156 showed approximately 30% cell death at a 300 μM concentration.

However, NCCPF 470157 was not responsive (Supplementary Figure 2D).

### *C. albicans* can develop resistance to F1853-0039

Finally, we queried whether *C. albicans* can become resistant to F11853-0039. Cells were grown in the presence of 1 μM F1853-0039 for 2 weeks as explained in Materials and Methods, and MIC_50_ was subsequently calculated. We found that the cells indeed become resistant to F1853-0039; however, the mechanism of resistance needs to be further deciphered (Supplementary Fig. 3B).

## DISCUSSION

*C. albicans* is a commensal pathogen that is fatal to immunocompromised patients. The ability of the pathogen to switch from yeast to hyphae, as well as contact sensing and secretion of hydrolases, underlies its pathogenicity [42]. Limited drug options are available at our disposal to counter infection cases, and drug resistance is a severe concern [43]. Therefore, searching for new drug targets and potential pathogen inhibitors is necessary.

The epigenetic modulators comprising ATP-dependent chromatin remodelling proteins and histone modifying enzymes regulate gene expression and DNA repair processes [44,45]. Thus, these enzymes have emerged as drug targets for both infectious and non-infectious diseases. For example, histone deacetylases of *C. albicans* regulate yeast-to-hyphae transition, white-to-opaque transition, and biofilm formation. Inhibitors against histone deacetylase, when used in combination with existing antifungal agents, show enhanced anti-Candida effects [3].

Previously, we had shown that *C. albicans* Fun30, an ATP-dependent chromatin remodelling protein, regulates the expression of RTT109, a histone acetyltransferase, present in the organism. We have also shown that *FUN30 i*s not essential for the viability of *C. albicans* but is required for the DNA damage response, as it not only mediates end resection but also co-regulates the expression of DNA damage response genes [15]. We, therefore, hypothesised that *Ca*Fun30 might be a promising drug target.

Our current study provides proof-of-concept. Using in silico methods, we have identified ten potential inhibitors of *Ca*Fun30. *In vitro* binding studies showed that F1853-0039 binds to *Ca*Fun30 with a K_d_ in the sub-micromolar range and was able to significantly inhibit the ATPase activity of the protein. The IC_50_ was determined to be 15.8 μM against BWP17 and 19.1 μM against SN152, suggesting that the molecule is equally effective against both *C. albicans* strains. It needs to be noted that the molecule identified as an inhibitor with high affinity by the *in silico* method did not correspondingly show high affinity in the *in vitro* analysis. This is expected as the protein structure was predicted using homology modelling, and *in silico* analysis does not always corroborate with *in vitro* and *in vivo* experimental data. *In silico* ADME studies showed that the gastrointestinal tract does not absorb the molecule, and interestingly, *in vitro* MTT assay showed that the F1853-0039 is not toxic to THP-1 cells, suggesting that the molecule might have specificity towards the fungus. Co-culture studies confirmed that hyphae formation and *C. albicans* growth were inhibited by F1853-0039. Gene expression analysis showed that *ECE1* was downregulated and *NRG1*, encoding for a repressor of hypha formation, was upregulated. These changes in gene expression possibly contribute to hyphae inhibition. Further transcriptomic studies will be needed to understand the global changes in gene expression in response to F1853-0039.

Mechanistically, the small molecule inhibitor, F1853-0039, inhibits the ATPase activity of Fun30, leading to downregulation of chitin synthesis genes and alteration in expression of hyphal genes. Together, the cell wall composition is altered, and hyphae formation is blocked leading to reduced infection.

The specificity of the inhibitor for *C. albicans* is potentially exciting. *Ca*Fun30 and human SMARCAD1 share only 39.5% identity, even though the two proteins can functionally complement[15]. The specificity of the inhibitor towards *C. albicans* could be due to multiple reasons. F1853-0039 may have a higher affinity for *Ca*Fun30 compared to human SMARCAD1. Also, F1853-0039 probably has better permeability for fungal cells than human cells. Another key difference could lie in specific targets such that the inhibitor blocks critical pathways in *C. albicans* but not human cells. For example, the histone acetyltransferase, Rtt109, is unique to fungal systems and is required for GPI anchor biosynthesis and virulence [17,18]. *RTT109* expression is regulated by *Ca*Fun30 [15] and is downregulated in the presence of F1853-0039. Further, we have observed that the chitin levels, as measured by CFW stain as well as qPCR, are lower in the treated *C. albicans* cells as compared to the untreated samples. As chitin is specific to the fungal cells, therefore, the inhibitor may be more effective against *C. albicans* than against mammalian cells. There might also be compensatory pathways existing in human cells for the loss of human SMARCAD1, while such pathways might be absent in the fungal cells. For example, the loss of the mammalian SWI/SNF complex can be compensated by EP400/TIP60 [46]. Finally, these studies have been performed using cell lines, which might not accurately reflect the human system.

The inhibitor shows synergistic action with the existing antifungal drugs, indicating that it can be used in combinatorial therapy. However, the pathogen also develops resistance to the molecule on prolonged exposure, suggesting that the compound is not infallible. Therefore, further studies need to be performed to evaluate the potential of F1853-0039 as an effective lead molecule.

The studies presented here provide an initial proof-of-concept that inhibitors can be developed against *Ca*Fun30. Additional studies, including animal models, testing against other *Candida* spp., and synthesis of better inhibitor molecules could help develop new lead molecules effective against Candidiasis.

## Author information

### Corresponding author

Rohini Muthuswami, Room # 306, School of Life Sciences, JNU, New Delhi-110067, India. Email

### Author Contributions

Conceptualisation, R.M.; Methodology, R.M., and K.P.; Investigation, K.P., P.P.R., F.K., S.S., R.V., and V.A.; Writing—original draft, R.M., S.G.N., and K.P.; Writing—review and editing, R.M., S.G.N., and K.P.; Funding acquisition, R.M., and S.G.N.; Supervision, R.M. and S.G.N. All authors have read and agreed to the published version of the manuscript.

## Funding

The work was supported by MHRD-STARS (STARS/APR2019/BS/224/FS). We acknowledge the facilities/laboratories supported by DBT BUILDER (BT/INF/22/SP45382/2022) and DST FIST-II (SR/FST/LSII-046/2016(C)). PPR acknowledges DBT for M.K.Bhan, YRF for fellowship.

### Data Availability Statement

The authors confirm that the data supporting the findings of this study are available within The article and its supplementary materials.

