## Supplementary figures and images for "Antifungal therapeutic potential of *Candida albicans* Fun30: screening and validation of novel inhibitors against *Candida albicans*"

### Supplementary Figure S1

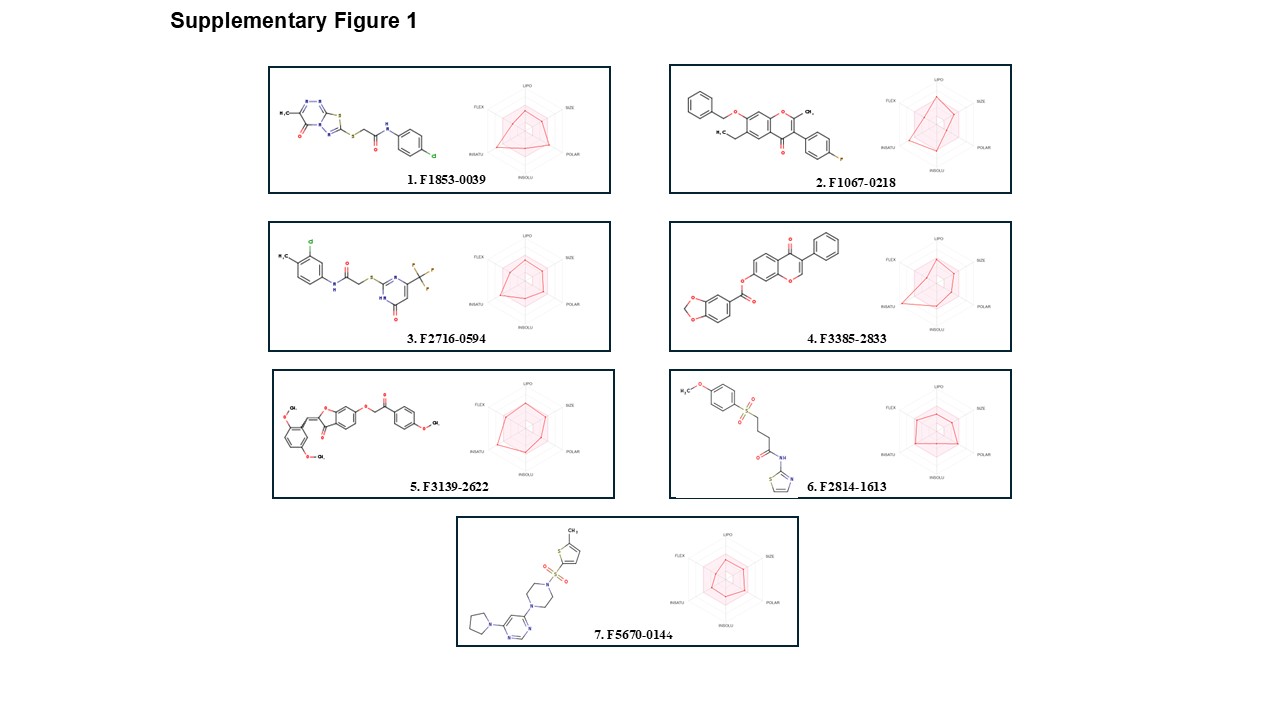

### Supplementary Figure S2

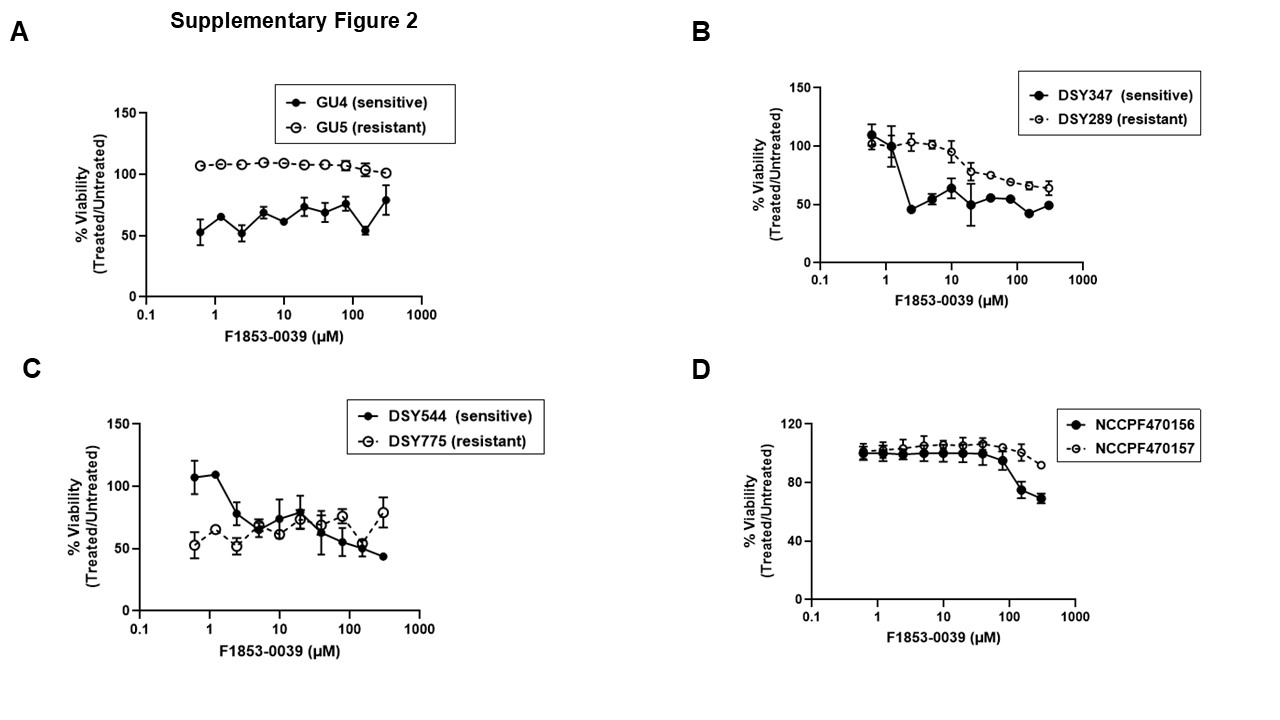

### Supplementary Figure S3

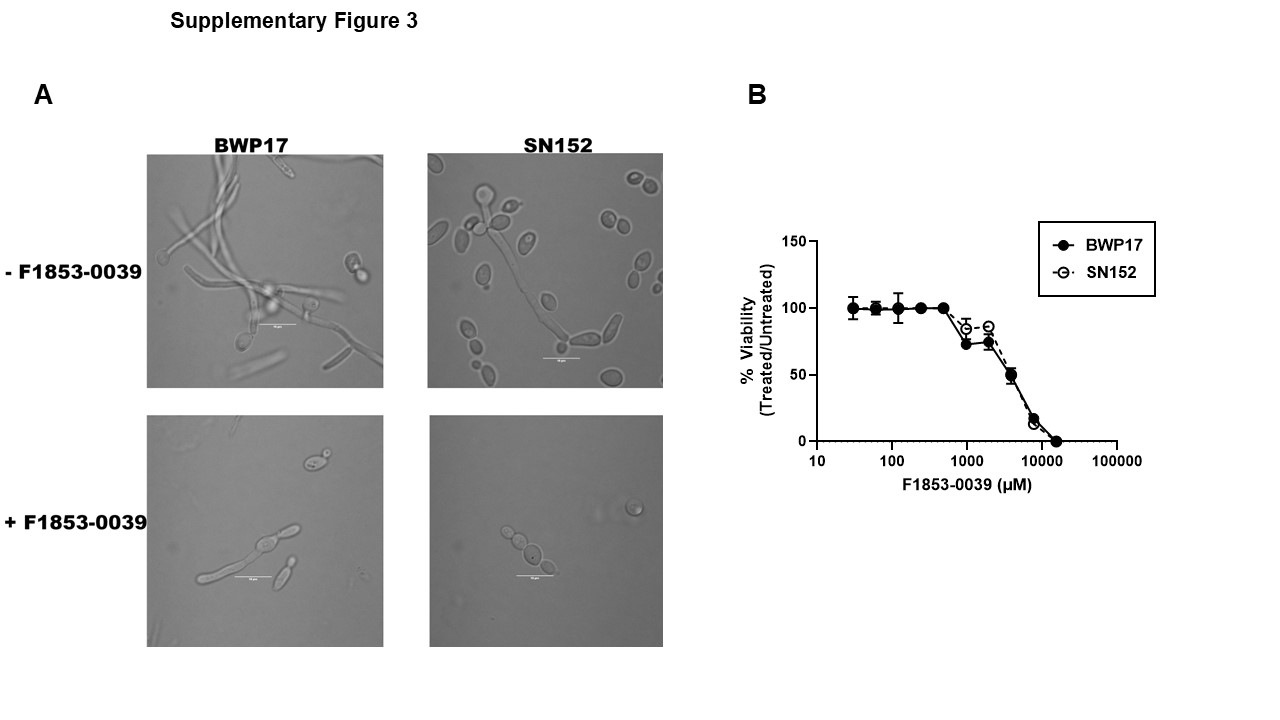
