## Supplementary File for "Antifungal therapeutic potential of *Candida albicans* Fun30: screening and validation of novel inhibitors against *Candida albicans*"

**SUPPLEMENTARY FIGURE LEGENDS**

**Supplementary Figure 1:** Structure of lead compounds and their bioavailability radar chart. The coloured zone shows the suitable physicochemical space for oral bioavailability. F1853-0039 shows significant polarity and solubility scores.

**Supplementary Figure 2:** Effect of F1853-0039 on pair clinical isolates of *Candida albicans* (A). GU4 (drug-sensitive) and GU5 (drug-resistant). (B). DSY 347 (drug-sensitive) and DSY 289 (drug-resistant). (C). DSY 544 (drug-sensitive) and DSY 775 (drug-resistant). (D). Effect of F1853-0039 on drug-resistant isolates of *Candida auris.* The data is presented as the average ± SD of two independent replicates.

**Supplementary Figure 3:** Effect of F1853-0039 on hyphae formation and development of resistance. (A). Hyphae formation was observed under confocal microscope in BWP17 and SN152 cells in the absence and presence of 100 μM inhibitor. (B) MIC of BWP17 and SN152 cells was calculated after growing the cells in 1 μM inhibitor for 10 days.

**Supplementary table 1: ADME analysis of lead compounds**

| Molecule | F1853-0039 | F1067-0218 | F2716-0594 | F3385-2833 | F3139-2622 | F2814-1613 | F5670-0144 |
| --- | --- | --- | --- | --- | --- | --- | --- |
| Canonical SMILES | CC1=NN=C2SC(SCC(=O)NC3=CC=C(Cl)C=C3)=NN2C1=O | CCC1=CC2=C(OC(C)=C(C2=O)C2=CC=C(F)C=C2)C=C1OCC1=CC=CC=C1 | CC1=C(Cl)C=C(NC(=O)CSC2=NC(=CC(=O)N2)C(F)(F)F)C=C1 | O=C(OC1=CC=C2C(=O)C(=COC2=C1)C1=CC=CC=C1)C1=CC=C2OCOC2=C1 | COC1=CC=C(C=C1)C(=O)COC1=CC=C2C(=O)\C(OC2=C1)=C\C1=C(OC)C=CC(OC)=C1 | COC1=CC=C(C=C1)S(=O)(=O)CCCC(=O)NC1=NC=CS1 | CC1=CC=C(S1)S(=O)(=O)N1CCN(CC1)C1=NC=NC(=C1)N1CCCC1 |
| Formula | C13H10ClN5O2S2 | C25H21FO3 | C14H11ClF3N3O2S | C23H14O6 | C26H22O7 | C14H16N2O4S2 | C17H23N5O2S2 |
| MW | 367.83 | 388.43 | 377.77 | 386.35 | 446.45 | 340.42 | 393.53 |
| #Heavy atoms | 23 | 29 | 24 | 29 | 33 | 22 | 26 |
| #Aromatic heavy atoms | 15 | 22 | 12 | 22 | 18 | 11 | 11 |
| Fraction Csp3 | 0.15 | 0.16 | 0.21 | 0.04 | 0.15 | 0.29 | 0.53 |
| #Rotatable bonds | 5 | 5 | 6 | 4 | 8 | 8 | 4 |
| #H-bond acceptors | 5 | 4 | 6 | 6 | 7 | 5 | 5 |
| #H-bond donors | 1 | 0 | 2 | 0 | 0 | 1 | 0 |
| MR | 90.46 | 113.6 | 85.39 | 105.38 | 121.5 | 85.15 | 113.58 |
| TPSA | 142.79 | 39.44 | 100.15 | 74.97 | 80.29 | 121.98 | 106.26 |
| iLOGP | 2.41 | 4.21 | 2.24 | 3.62 | 4.04 | 2.01 | 3.13 |
| XLOGP3 | 2.75 | 5.92 | 3.01 | 4.4 | 4.89 | 1.66 | 2.64 |
| WLOGP | 2.05 | 6.32 | 4.44 | 4.41 | 4.48 | 3.23 | 1.9 |
| MLOGP | 2.13 | 4.08 | 2.63 | 2.75 | 1.46 | 0.88 | 1.31 |
| Silicos-IT Log P | 2.77 | 7.29 | 4.34 | 4.82 | 5.36 | 2.68 | 1.68 |
| Consensus Log P | 2.42 | 5.56 | 3.33 | 4 | 4.05 | 2.09 | 2.13 |
| ESOL Log S | -4.01 | -6.21 | -4.05 | -5.3 | -5.56 | -2.84 | -3.99 |
| ESOL Solubility (mg/ml) | 3.63E-02 | 2.40E-04 | 3.35E-02 | 1.92E-03 | 1.22E-03 | 4.94E-01 | 4.01E-02 |
| ESOL Solubility (mol/l) | 9.87E-05 | 6.18E-07 | 8.86E-05 | 4.96E-06 | 2.73E-06 | 1.45E-03 | 1.02E-04 |
| ESOL Class | Moderately soluble | Poorly soluble | Moderately soluble | Moderately soluble | Moderately soluble | Soluble | Soluble |
| Ali Log S | -5.4 | -6.52 | -4.78 | -5.69 | -6.31 | -3.84 | -4.52 |
| Ali Solubility (mg/ml) | 1.45E-03 | 1.17E-04 | 6.30E-03 | 7.86E-04 | 2.18E-04 | 4.97E-02 | 1.18E-02 |
| Ali Solubility (mol/l) | 3.95E-06 | 3.00E-07 | 1.67E-05 | 2.03E-06 | 4.88E-07 | 1.46E-04 | 3.00E-05 |
| Ali Class | Moderately soluble | Poorly soluble | Moderately soluble | Moderately soluble | Poorly soluble | Soluble | Moderately soluble |
| Silicos-IT LogSw | -4.92 | -10.18 | -6.42 | -8.02 | -8 | -5.05 | -3.91 |
| Silicos-IT Solubility (mg/ml) | 4.43E-03 | 2.57E-08 | 1.43E-04 | 3.69E-06 | 4.49E-06 | 3.06E-03 | 4.89E-02 |
| Silicos-IT Solubility (mol/l) | 1.21E-05 | 6.61E-11 | 3.78E-07 | 9.55E-09 | 1.00E-08 | 8.99E-06 | 1.24E-04 |
| Silicos-IT class | Moderately soluble | Insoluble | Poorly soluble | Poorly soluble | Poorly soluble | Moderately soluble | Soluble |
| GI absorption | Low | High | High | High | High | High | High |
| BBB permeant | No | No | No | No | No | No | No |
| Pgp substrate | No | Yes | No | No | No | No | Yes |
| CYP1A2 inhibitor | Yes | Yes | Yes | Yes | No | No | No |
| CYP2C19 inhibitor | Yes | Yes | Yes | Yes | Yes | Yes | Yes |
| CYP2C9 inhibitor | Yes | No | No | Yes | Yes | Yes | Yes |
| CYP2D6 inhibitor | No | No | No | No | No | No | Yes |
| CYP3A4 inhibitor | No | Yes | No | Yes | Yes | No | Yes |
| log Kp (cm/s) | -6.59 | -4.47 | -6.47 | -5.53 | -5.55 | -7.2 | -6.83 |
| Lipinski #violations | 0 | 0 | 0 | 0 | 0 | 0 | 0 |
| Ghose #violations | 0 | 1 | 0 | 0 | 0 | 0 | 0 |
| Veber #violations | 1 | 0 | 0 | 0 | 0 | 0 | 0 |
| Egan #violations | 1 | 1 | 0 | 0 | 0 | 0 | 0 |
| Muegge #violations | 0 | 1 | 0 | 0 | 0 | 0 | 0 |
| Bioavailability Score | 0.55 | 0.55 | 0.55 | 0.55 | 0.55 | 0.55 | 0.55 |
| PAINS #alerts | 0 | 0 | 0 | 0 | 0 | 0 | 0 |
| Brenk #alerts | 0 | 0 | 0 | 1 | 1 | 0 | 0 |
| Leadlikeness #violations | 1 | 2 | 1 | 2 | 3 | 1 | 1 |
| Synthetic Accessibility | 3.06 | 3.59 | 2.54 | 3.52 | 3.96 | 2.94 | 3.5 |

**Supplementary Table 2.** List of primers used for qPCR

| **Transcript name** | **Forward Primer (5' →3')** | **Reverse Primer (5' →3')** |
| --- | --- | --- |
| *RDN18* | CGGCACCTTACGAGAAATCA | CCACCACCCACAAAATCAA |
| *FUN30* | GGATTGGCAGCTAGATTCAG | TCGCTTTGGTACTAGATGGT |
| *GPI15* | CCACGATTATGGGCAGGTTA | CGCGGTAAGAATTCTGGAAA |
| *Rtt109* | TCGTTGATTGGATGCTGTAAGG | ACCAGCTTCAACAGGTTCATAA |
| *CHS1* | GAGATTAGTTGGTGGCAAAGCAG | CCGTATTTGGCAGCTCTTAAGG |
| *CHS2* | GTTGTCATTGCCTTGGTTTTAG | GCATGTTCACTTGCCTTAATTTC |
| *HWP1* | GATCCCAGATATCCCAGAAAAG | GTAGCTGGAGTAGTTGATTCTG |
| *ECE1* | GTCAGATTGCCAGAAATTGTTG | GATCTAGTAATGAGTTGTGGGAG |
| *BCR1* | GTATAACTTCCAACCTCCATATCC | GCCATAGGATCAAAAGTGGTAT |
| *ADH5* | CTCCTTACCATGCTATCAAGTC | GCTGTCTGGTAATTCACTGTAA |
| *CSH1* | CAATTCATCTCCATGCAAAGTC | TAATCTTATCTGCGTCTCTGAC |
| *FLO8* | CAGATATGGAGACGGTAACAGTA | GTTGATTTTGCATACCACTAGC |
| *NRG1* | GGACATTTAGCTAGACACAATC | CATGTGAAGCTTCTAAAGTCC |

**Supplementary Table 3: qPCR conditions**

| 95^o^C | 20 sec | Step 1 | X1 | Initial denaturation |
| --- | --- | --- | --- | --- |
| 95^o^C | 3 sec | Step 2 | X40 | Cycling stage |
| 60^o^C | 30 sec | Step 3 | X40 | Cycling stage |
| 95^o^C | 15 sec | Step 4 | X1 | Melt curve stage |
| 60^o^C | 1 min | Step 5 | X1 | Melt curve stage |
| 95^o^C | 15 sec | Step 6 | X1 | Melt curve stage |
| 60^o^C | 15 sec | Step 7 | X1 | Melt curve stage |
| 4^o^C | hold | Step 8 | X1 | hold |
